# Argon plasma-modified bacterial cellulose filters for protection against respiratory pathogens

**DOI:** 10.1101/2022.04.28.489859

**Authors:** Anna Żywicka, Daria Ciecholewska-Juśko, Magdalena Szymańska, Radosław Drozd, Peter Sobolewski, Adam Junka, Selestina Gorgieva, Miroslawa El Fray, Karol Fijałkowski

## Abstract

Due to the global spread of the SARS-CoV-2 virus and the resultant pandemic, there has been a major surge in the demand for surgical masks, respirators, and other air filtration devices. Unfortunately, the fact that these filters are made of petrochemical-derived, non-biodegradable polymers means that the surge in production has also led to a surge in plastic waste. In this work, we present novel, sustainable filters based on bacterial cellulose (BC) functionalized with low-pressure argon plasma (LPP-Ar). The “green” production process involved BC biosynthesis by *Komagataeibacter xylinus*, followed by simple purification, homogenization, lyophilization, and finally LPP-Ar treatment. The obtained LPP-Ar-functionalized BC-based material (LPP-Ar-BC-bM) showed excellent antimicrobial and antiviral properties, with no cytotoxicity versus murine fibroblasts *in vitro*. Further, filters consisting of three layers of LPP-Ar-BC-bM had >99% bacterial and viral filtration efficiency, while maintaining sufficiently low airflow resistance (6 mbar at an airflow of 95 L/min). Finally, as a proof-of-concept, we were able to prepare 80 masks with LPP-Ar-BC-bM filter and ~85% of volunteer medical staff assessed them as good or very good in terms of comfort. We conclude that our novel sustainable, biobased, biodegradable filters are suitable for respiratory personal protective equipment (PPE), such as surgical masks and respirators. Further, with scale-up, they may be adapted for indoor air handling filtration in hospitals or schools.

**Graphical abstract:** 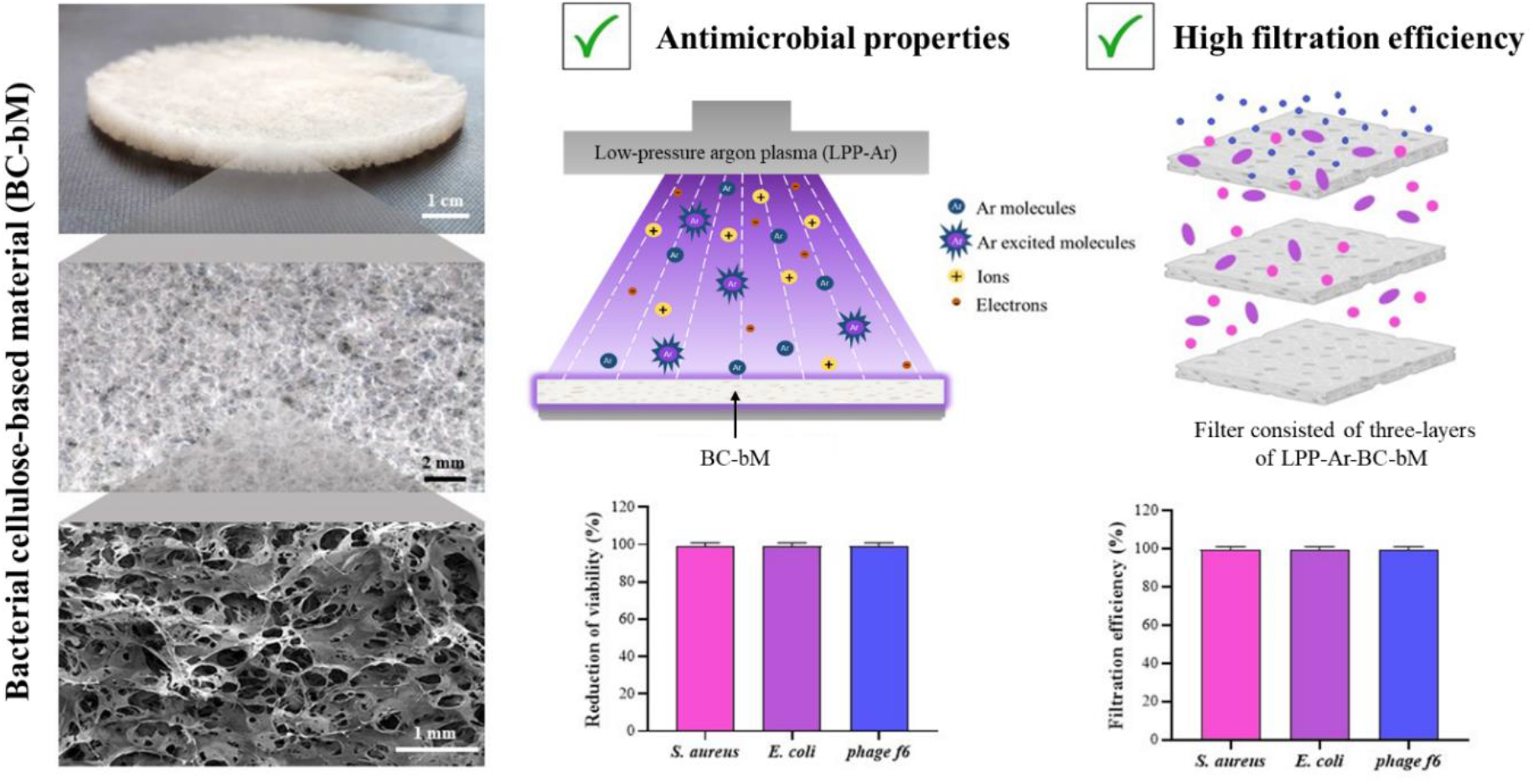

## Introduction

Due to the global spread of the SARS-CoV-2 virus and the resultant pandemic, there has been a major surge in the demand for surgical masks, respirators, and other air filtration devices. Not surprisingly, this has led to a significant increase in their industrial production^1^. Importantly, the SARS-CoV-2 pandemic has also led to a renewed recognition of the danger posed by airborne transmission of respiratory pathogens^2^, meaning that the demand for air filtration materials, whether for personal protective equipment (PPE) or indoor air handling will remain high. In this context, the primary focus is on particle filters that trap solid and liquid aerosols, particularly those containing biological contaminants (fungi, bacteria, viruses), although filters that protect against both gaseous and particulate contaminants are also possible^3^. Depending on the type of filter and the properties of the material used, the level of particle filtration efficiency (PFE), bacterial filtration efficiency (BFE), or viral filtration efficiency (BFE) may differ significantly^4,5^. Currently, the majority of filters use fibers composed of polypropylene (PP), poly(ethylene terephthalate)(PET), poly(tetrafluoroethylene), (PTFE), polyamide (PA), or polycarbonate (PC)^6^. As a result, the surge in production of air filters sparked by the SARS-CoV-2 pandemic has a large carbon footprint, while also generating a surge in non-degradable plastic waste^1,7^ –all while the world is already grappling with the environmental impact of millions of tons of existing non-degradable plastic waste^8^.

Therefore, there is a clear and urgent need to develop novel filter materials with sustainability in mind, with the ultimate goal of facilitating a circular economy in this product area. Ideally, such filters would consist of biodegradable materials that can be obtained sustainably, from renewable resources. One example of more sustainable filter material is plant cellulose. However, to increase filtration efficiency, plant cellulose is frequently modified using synthetic polymers e.g. poly(ethyleneimine) and cellulose derivatives, such as cellulose nitrate, mixtures of cellulose esters, cellulose triacetate, or cellulose acetate phthalate^6,9–12^. Additionally, while it is renewable and biodegradable, the overall life cycle of plant cellulose has a high global warming impact plus added drawbacks, including pesticide use and high arable land and water requirements^13^. Biotechnology offers a promising alternative: bacterial cellulose (BC), a biopolymer produced by a spectrum of aerobic, non-pathogenic, Gram-negative bacteria, such as *Komagataeibacter xylinus*. Importantly, BC consists of loosely arranged, highly crystalline fibrils ~100 nm in diameter and is characterized by high mechanical strength, chemical stability, biocompatibility, and biodegradability^14–16^. Combined these aspects make BC well suited as an alternative to synthetic polymers used as filtration membranes.

Filtration efficiency is the most important parameter in terms of air filter usability and helps determine potential applications. However, it is important to note that pathogens that are just trapped physically within a filter can still pose a biological hazard, because of the risk that they could be released back into the airstream. As a result, many studies have been devoted to fabricating air filters that not only sequester pathogens but also kill them. For example, filters have been proposed that incorporate antimicrobial agents such as ε-polylysine and natamycin, silver and zinc nanoparticles, or plant extracts from eucalyptus, grapefruit, or propolis^15,17–20^. Likewise, several patents describe BC-based air filters that include antimicrobial components, such as silver^21^, zeolite-supported silver nanoparticles^22^, or chitosan^23^. However, the emergence and spread of microbes resistant to chemical agents^24^, combined with the need to improve sustainability, motivate further research and development into greener BC surface modification strategies.

One alternative approach involves the use of low-temperature plasma (LTP, also known as *cold plasma*), a partially ionized gas with a variety of electrons, ions, free radicals, and excited atoms and molecules^25^. The interactions of the plasma with the surface of polymeric materials can result in major improvements in wettability and surface charge, changes in surface topography, and can impart antimicrobial activity^26–28^. LTP treatment can be performed at atmospheric or low pressure (APP, LPP, respectively). APP treatment requires the application of higher voltages for gas breakdown and tends to result in more heating, due to enhanced collisions between electrons and gas molecules^29^. Typically, APP modification is carried out using a nozzle system and to treat large surfaces, a large number of nozzles is required. In contrast, LPP is carried out in vacuum chambers which limit the size that can be modified at one time but improves the plasma distribution, ensuring more even exposure of the surface of the sample^30^. Therefore, the LPP is used commonly for biomedical applications^26,31^. Importantly, LPP processes do not require any solvents and do not generate any waste, making them inherently “green”^32^.

The LPP process can use several different inert or reactive gases, which affect the relative amounts and types of surface functional groups formed, such as –OH, –CHO, –COOH, or – NH_2_. Among the inert gases, argon is the most common^30^, because it has good thermal conductivity, a low electron affinity, a relatively high mass, low ionization energy, and a low rate of electrode erosion^33^. Importantly, several studies have shown that LPP modification using argon plasma (LPP-Ar) can improve the antibacterial properties of polymers. For example, polyester fabrics treated with LPP-Ar had improved activity against both Gram-positive (*S. aureus*) and Gram-negative (*E. coli*) bacteria^34^. Likewise, Slepicka et al. showed that LPP-Ar can improve the antibacterial properties of polytetrafluoroethylene (PTFE) nanotextiles and tetrafluoroethylene-perfluoro(alkoxy vinyl ether) (PFA) films^35^. However, to our knowledge, there are no reports on the use of LPP-Ar to improve the antimicrobial properties of BC for use as air filters.

Our goal in this work was to develop fully biobased BC-based filters, displaying excellent bacterial and viral filtration efficiency and adequate airflow resistance parameters, as well as having antibacterial and antiviral properties. Towards this aim, we developed a homogenization and lyophilization procedure for obtaining BC-based material (BC-bM) from BC pellicles. Next, we showed for the first time that LPP-Ar treatment of the BC-bM results in improved antibacterial and antiviral properties. We demonstrated that the developed LPP-Ar-functionalized BC-bM (LPP-Ar-BC-bM) meets the requirements described in the relevant standards including the European standard *EN 14683 + AC:2019-09 Medical masks – Requirements and test methods*, EN 13274-3:2008 *Respiratory Protective Devices – Methods of Test – Part 3: Determination of Breathing Resistance* and ISO 10993-5:2009: *Biological evaluation of medical devices; Part 12: Sample preparation and reference materials* and *Part 5: Tests for in vitro cytotoxicity*. Finally, as a proof-of-concept, we were able to prepare 80 masks with LPP-Ar-BC-bM filter for user experience testing by volunteer medical staff. Ultimately, the developed filters could play a valuable role as part of protective measures during the current SARS-CoV-2 and any future airborne pandemics.

## Results

### Optimization of BC-bM preparation process

The first step of our work was to develop a process for converting BC pellicles obtained from *K. xylinus* cultures into BC-bM that had homogeneous macrostructure and sufficiently low airflow resistance to be usable as filters in face masks and other respiratory PPE, based on EN 13274-3:2008. In preliminary work, we tested native BC that was only subjected to purification and dehydration (dried at 60 °C or lyophilized). However, it was not possible to measure the airflow resistance of dry BC (dried at 60 °C), because the pellicles were too brittle and ruptured during testing. Similarly, lyophilized native BC did not provide sufficient porosity and air permeability (Table 1, Fig. 1). These results led us to develop a two-step homogenization and lyophilization process that yielded three-dimensional, sponge-like BC-bMs (Fig. 1, Supplementary Fig. 1).

**Table 1.** The airflow resistance of BC pellicles and BC-bMs.

|  | Type of sample | Airflow resistance [mbar] |  |
| --- | --- | --- | --- |
|  |  | Set flow: 35 L/min | Set flow: 95 L/min |
| BC pellicles | Dry | not measurable |  |
|  | Freeze-dried | 4.6±0.43* | 12.5±1.38* |
| BC-bMs | BC pulp dilution 2:1 | 4.3±0.37* | 10.9±1.49* |
|  | BC pulp dilution 1:2 | 0.60±0.35 | 2.4±0.65 |
|  | **BC pulp dilution 1:1** | **1.0±0.03** | **4.0±0.32** |

Data were presented as a mean ± standard error of the mean (SEM). \* Results exceeding the limit values of air resistance for P1, P2, or P3 class filters according to European standards EN 13274-3:2008. Data in bold were selected as optimal for process parameters.

We optimized the production process of BC-bMs in terms of airflow resistance, the key parameter for use in respiratory PPE. We tested the effect of process parameters: pulp volume, pulp density, and freezing temperature. A summary of the influence of these parameters on airflow resistance is presented in Table 1, while Fig. 1 presents representative micrographs showing macroscopic homogeneity and porosity of representative BC-bMs. Based on airflow resistance measurements and macro-morphological appearance, we selected the following process parameters as optimal: 1:1 mass ratio of BC pulp to water, a volume of 80 mL (in a 144 cm^2^ square petri dish), and freezing temperature of −18 °C (indicated in bold in Table 1 and in green on Fig. 1). Using lower pulp density, lower pulp volume, or lower temperature resulted in materials that met airflow resistance requirements but had irregular and heterogeneous macromorphology (Table 1, Fig. 1). On the other hand, higher pulp density or larger volumes resulted in materials with high airflow resistance that exceeded levels specified by EN 13274-3:2008 standard (Table 1, Fig. 1).

### Optimization of BC-bM functionalization process

Once we established and optimized the process for obtaining homogenous BC-bM that fulfilled the requirements of EN 13274-3:2008 in terms of airflow resistance, we proceeded to develop an LPP-Ar functionalization process aimed at improving antibacterial and antiviral properties. We varied the treatment time from 1 to 30 min, to optimize the LPP-Ar functionalization process.

First, we used ATR-FTIR to assess changes in the chemical composition of LPP-Ar-BC-bMs depending on the duration of the functionalization process. The ATR-FTIR spectrum of BC-bM served as a control (Fig. 2a). In the first part of the spectrum (from 2600 cm^−1^ to 3600 cm^−1^), characteristic bands for OH of hydrogen bonds at 3334 cm^−1^ and CH at 2895 cm^−1^ were observed. The second region of spectra (from 1800 cm^−1^ to 800 cm^−1^) showed signals from adsorbed water at wavenumber 1648 cm^−1^, CH_2_ of C-6 at 1427 cm^1^, COH in-plane from C-2 and C-3 at 1134 cm^−1^, OC of β-glycosyl linkage at 1161 cm^1^, CO at C-6 at 1030 cm^−1^ and COC of β-glycosylic linkage characteristic for amorphous BC fraction at 897 cm^−1^.

**Fig. 2.**
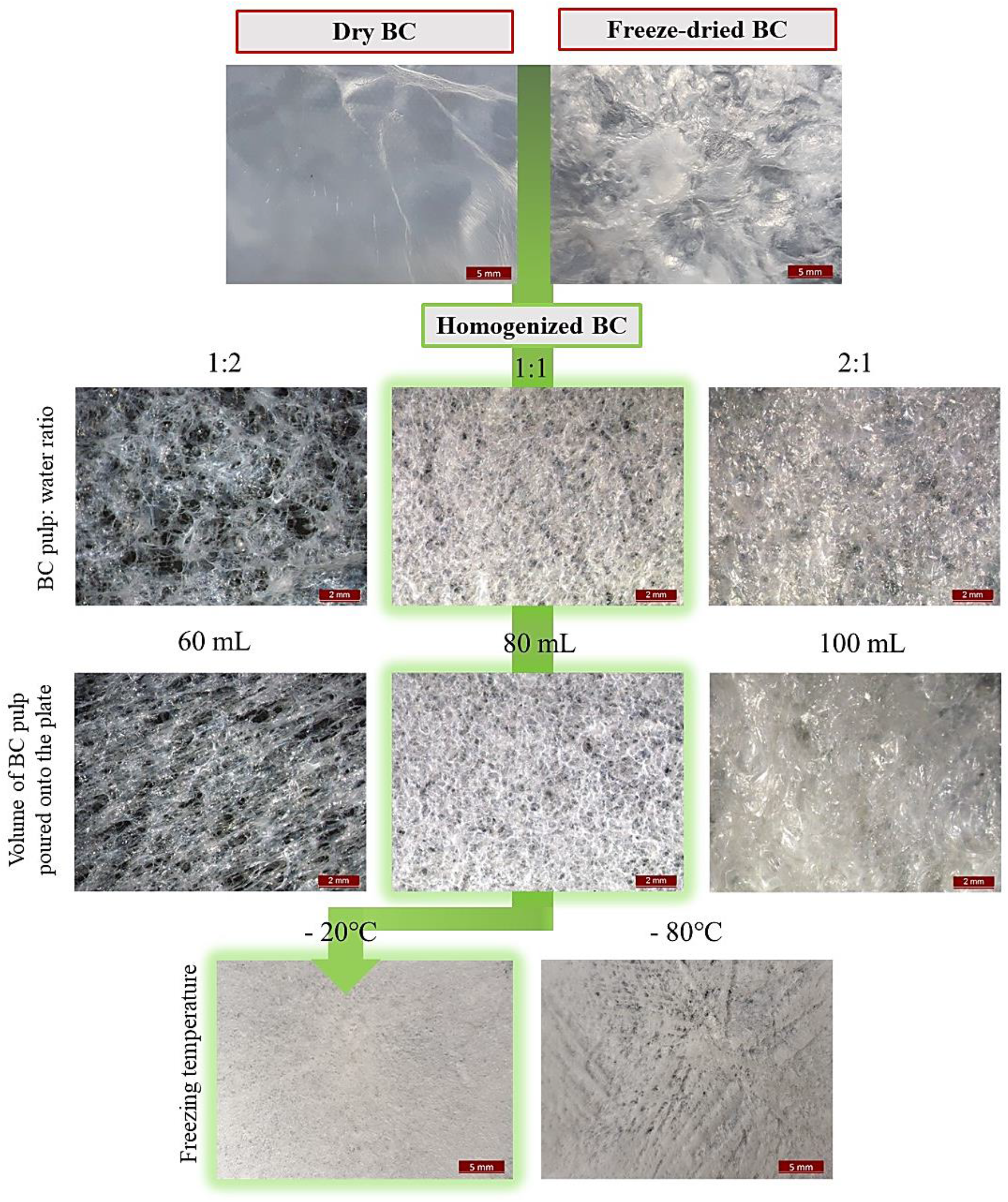
Representative micrographs showing the macro-morphological differences in BC pellicles and BC-bMs depending on different process parameters. Images were taken using stereoscopic microscope.

Following functionalization, the spectra of the LPP-Ar-BC-bMs showed a time-dependent increase in the absorbance band at 1720 cm^−1^, which can be assigned to the carbonyl groups of the carboxyl group (Fig. 2a). The maximum absorption for this band was observed after 10 and 30 min of LPP-Ar treatment (Fig. 2b). At the same time, we did not observe significant changes in the spectral range 1160 cm^−1^ to 1057 cm^−1^ assigned to C-O-C asymmetric stretching and C-O stretching vibrations of glycosidic bonds of the cellulose backbone, indicating that LPP-Ar treatment did not result in significant degradation/depolymerization. However, we did observe that, after 30 min of LPP-Ar treatment, the samples became yellowish and more brittle, likely a heating effect (Supplementary Fig. 2). As a result, we selected 10 min as the optimal LPP-Ar treatment time.

2D ATR-FTIR spectra confirmed significant changes in the structure of LPP-Ar-BC-bM as a result of the functionalization process lasting for 10 min (Fig. 3c,d). Significant auto-bands were observed in the range of 987 cm^−1^, 1007 cm^−1^, 1035 cm^−1^, 1061 cm^−1^, 1088 cm^−1^, 1110 cm^−1^, 1320 cm^−1^, 1336 cm^−1^, 1427 cm^−1^, 1640 cm^−1^, and 1720 cm^−1^. The presence of intense auto-bands in the range of 980–1100 cm^−1^ may indicate the effect of LPP-Ar treatment on primary alcohols, mainly C6OH, present in the structure of D-glucopyranose subunits. LPP-Ar treatment may lead to their oxidation to aldehydes and carboxylic acids, which was confirmed by the presence of auto-bands at 1720 cm^−1^, as well as intense negative crossbands at 1720 cm^−1^ vs 987 cm^−1^, 1007 cm^−1^, 1035 cm^−1^, 1061 cm^−1^, 1088 cm^−1^, and 1110 cm^−1^. Importantly, the R-OH groups of D-glucopyranose units form a network of hydrogen bonds, stabilizing the crystal structure of BC cellulose. Thus, a decrease in crystallinity after 10 min of LPP-Ar treatment would be expected (1427 cm^−1^), as confirmed by auto-bands at 1320 cm^−1^, 1336 cm^−1^, 1427 cm^−1^ and negative cross-bands in the synchronous spectra at: 1720 cm^−1^ vs 1320 cm^−1^, 1336 cm^−1^.

**Fig. 3.**
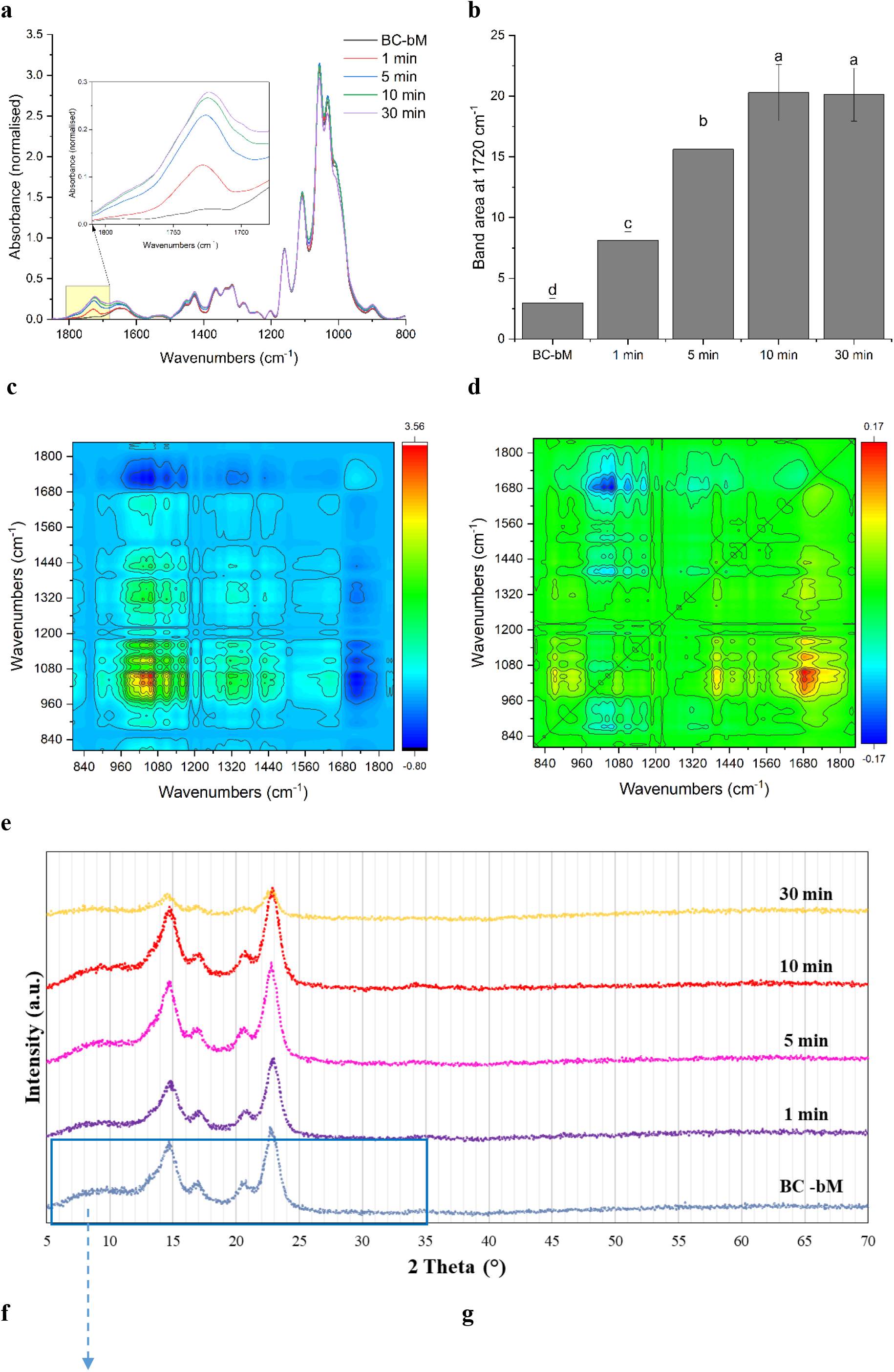
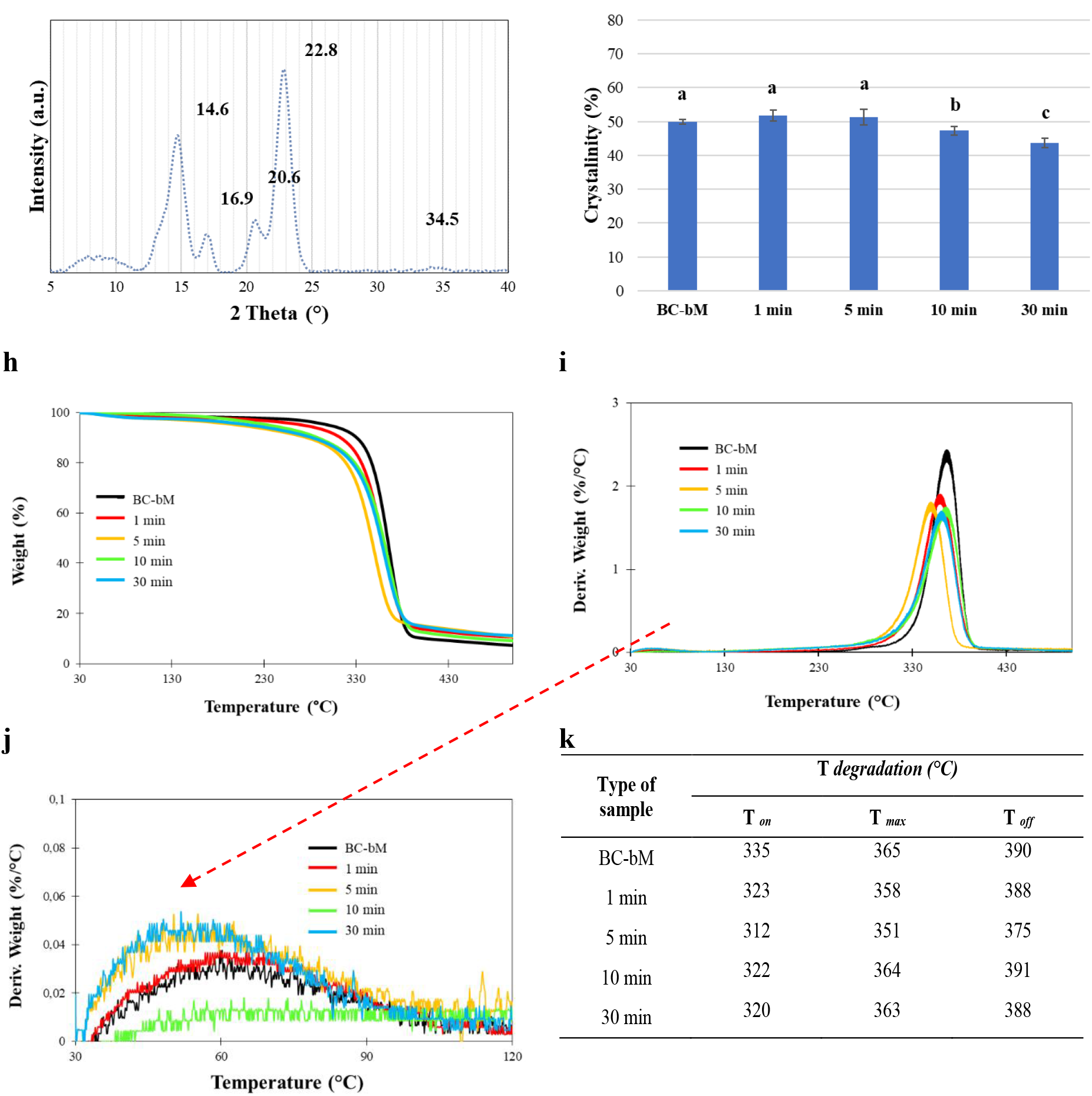
Chemical composition, crystallinity and thermal stability of BC-bM depending on the LPP-Ar treatment time. ATR-FTIR analysis of LPP-Ar-BC-bMs: **a)** region from 1850 cm^−1^ to 800 cm^−1^ depending on the duration of LPP-Ar treatment; **b)** differences in the area of the absorption band at 1720 cm^−1^ as a function of LPP-Ar treatment time; **c)** synchronous and **d)** asynchronous spectra from 2DCorr analysis of ATR-FTIR absorption spectra after 10 min of LPP-Ar treatment. XRD analysis of LPP-Ar-BC-bMs: **e)** full X-ray patterns of LPP-Ar-BC-bMs depending on the duration of LPP-Ar treatment; **f)** extracted area of interest with typical peaks related to BC crystallinity; **g)** crystallinity (%). Thermogravimetric analysis of LPP-Ar-BC-bMs: **h)** TGA curves, **i)** 1^st^ derivative curves; **j)** Magnified region of interest within 1^st^ derivative curves, related to water evaporation. **k)** Table insert with extracted data for T*_on_*, T*_max_*, and T*_off_* temperatures from the thermal degradation curves. Data in **b)** and **g)** were presented as a mean ± standard error of the mean (SEM). Different letters indicate statistically significant differences (p<0.05).

We repeated the ATR-FTIR measurements after 1, 2, and 3 months of storage of LPP-Ar-BC-bMs at room temperature in a desiccator. We did not observe any significant reduction in the intensity of the absorption band at 1720 cm^−1^ during storage, as compared to the initial value immediately after LPP-Ar treatment (Supplementary Fig. 3), indicating that the obtained materials have a shelf-life of at least 3 months.

To further examine the effect of the functionalization process on the crystallinity of the LPP-Ar-BC-bMs, we used XRD analysis. Overall, all of the complete XRD spectral lines were similar, with peaks typical of cellulose I structure (Fig. 3e). The four characteristic crystalline peaks for cellulose I were observed at 14.6°, 16.9°, 20.6°, and 22.8° (Fig. 3f). The XRD data indicated that 1 and 5 min of LPP-Ar treatment had no impact on the crystallinity, which was ~50%. However, consistent with ATR-FTIR analysis, after 10 min of LPP-Ar treatment, the crystallinity of LPP-Ar-BC-bM decreased to ~47%, with a further decrease after 30 min (Fig. 3g).

To assess the influence of plasma treatment on thermal stability of LPP-Ar-BC-bMs, we used TGA analysis. TGA and derivative (DTA) curves presented in Fig. 3h, 3i, indicated changes in the behavior of LPP-Ar-BC-bMs depending on the duration of the functionalization process. The weight loss detected in region between 30 °C and 120 °C (Fig. 3j), is related to the evaporation of adsorbed water. As expected, LPP-Ar-BC-bMs, which would be expected to be more hydrophilic and have more adsorbed water, showed a trend towards greater mass loss in this region, as compared to the control BC-bM. The extracted data from the derivative curve, presented within table insert (Fig. 3k) imply on highest degradation onset temperature (T*_on_*) in BC-bM, indicating greater thermal stability, which may be related to higher crystallinity. On the other hand, the LPP-Ar-BC-bMs started to degrade at lower temperatures and resulted in higher residual (char) weight of ~10 %.

Based on the results obtained using ATR-FTIR, XRD, and TGA, it was concluded that 10 min was an optimal time for LPP-Ar treatment. Therefore, further analyses, presented below, were carried out only for LPP-Ar-BC-bMs functionalized for 10 min.

### Microstructure of LPP-Ar-BC-bM

In order to assess the effect of the functionalization process on the microstructure of the LPP-Ar-BC-bM functionalized for 10 min, we used SEM analysis. A clear effect of LPP-Ar treatment on the surface morphology (roughness and porosity) was observed (Fig. 4a, b, Supplementary Fig. 4). Both LPP-Ar-BC-bM and control BC-bM had differentiated pore structures, with both macropores (>100 μm) and micropores (<100 μm) visible. However, LPP-Ar-BC-bM was characterized by a higher degree of porosity compared to control BC-bM (Fig. 4a, b, Supplementary Fig. 4, Supplementary Fig. 5). The maximum, minimum, as well as average pore diameter of LPP-Ar-BC-bM were all significantly higher, as compared to the control BC-bM (Fig. 4c). At the same time, the variability in the size of the pores was also much greater in the case of LPP-Ar-BC-bM.

**Fig. 4.**
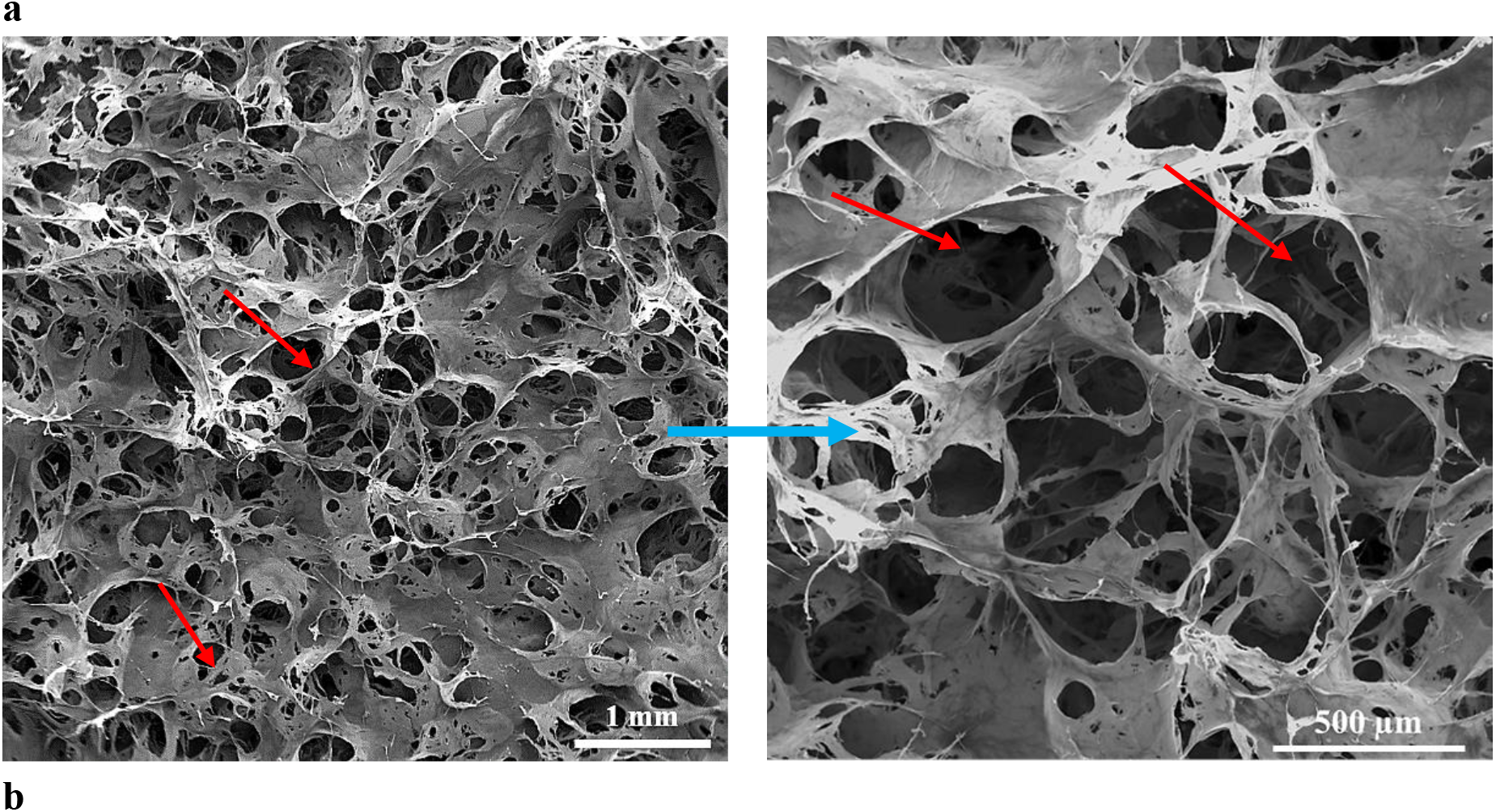
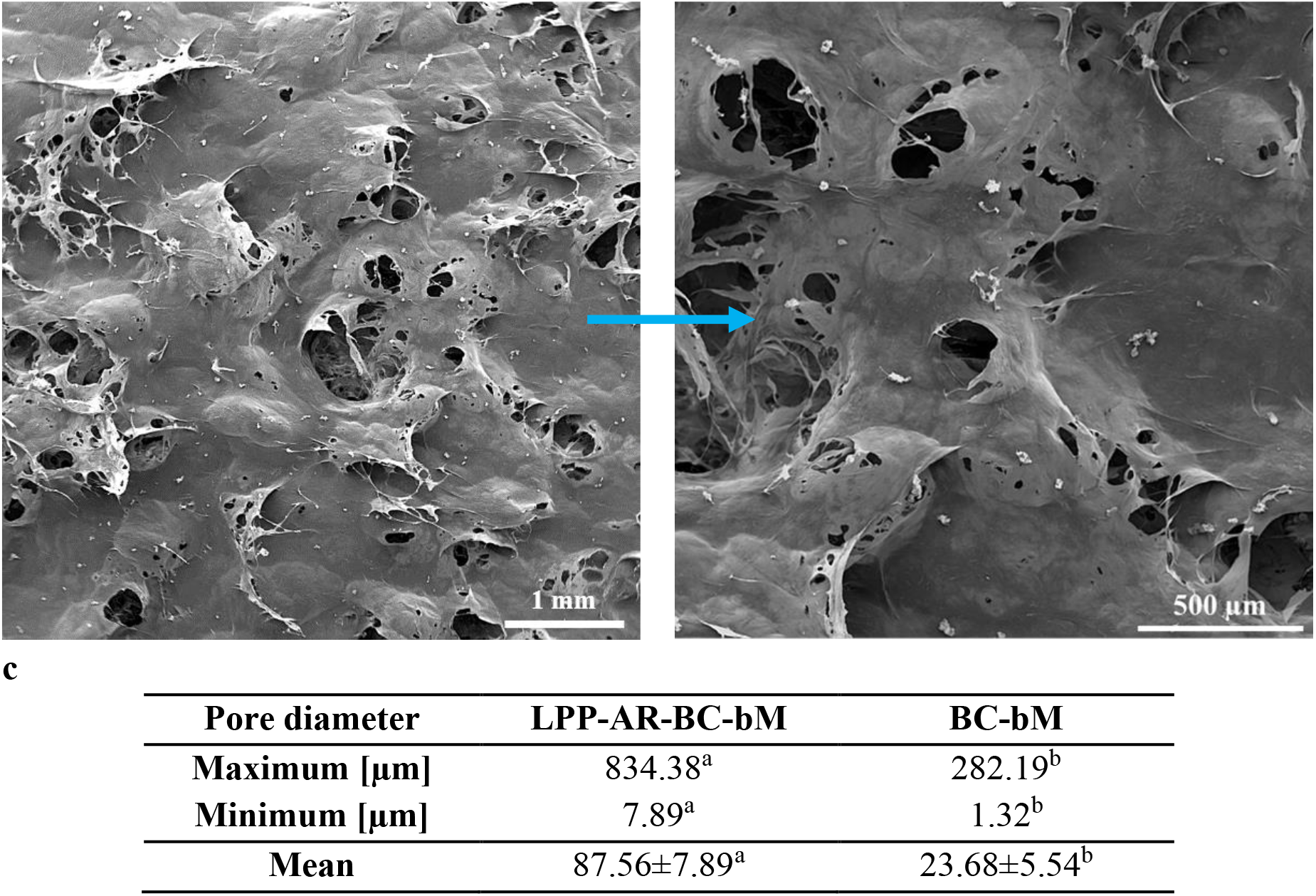
Microstructure of LPP-Ar-BC-bM. Representative SEM micrographs of **a)** LPP-Ar-BC-bM and **b)** BC-bM. **c)** Pore diameters of LPP-Ar-BC-bM v. BC-bM. Pores of diverse sizes visible on the surface of the sample **a)** and **b)** are marked with red arrows. Data in **c)** were presented as a mean ± standard error of the mean (SEM). Different letters indicate statistically significant differences (p<0.05).

### Antibacterial and antiviral properties of LPP-Ar-BC-bM

To test the hypothesis that LPP-Ar treatment would improve the antimicrobial and antiviral properties of our material, we performed tests in accordance with ISO 20743:2013 and ISO 18184:2019. Importantly, we confirmed that after 10 min of LPP-Ar treatment, LPP-Ar-BC-bM displayed strong antimicrobial activity against Gram-positive *S. aureus* and Gram-negative *E. coli* bacteria, as well as strong antiviral activity against bacteriophage Φ6 (Fig. 5a), an enveloped bacteriophage that can be used as a model surrogate for studying the surface and air survival of pathogenic viruses, including SARS-CoV-2^36^. In both cases, we observed a >99% reduction in cell/phage viability, as compared to the control BC-bM. Further, we repeated the tests with LPP-Ar-BC-bMs after 1, 2, and 3 months of storage and saw no reduction in antibacterial and antiviral activity (Fig. 5b).

**Fig. 5.**
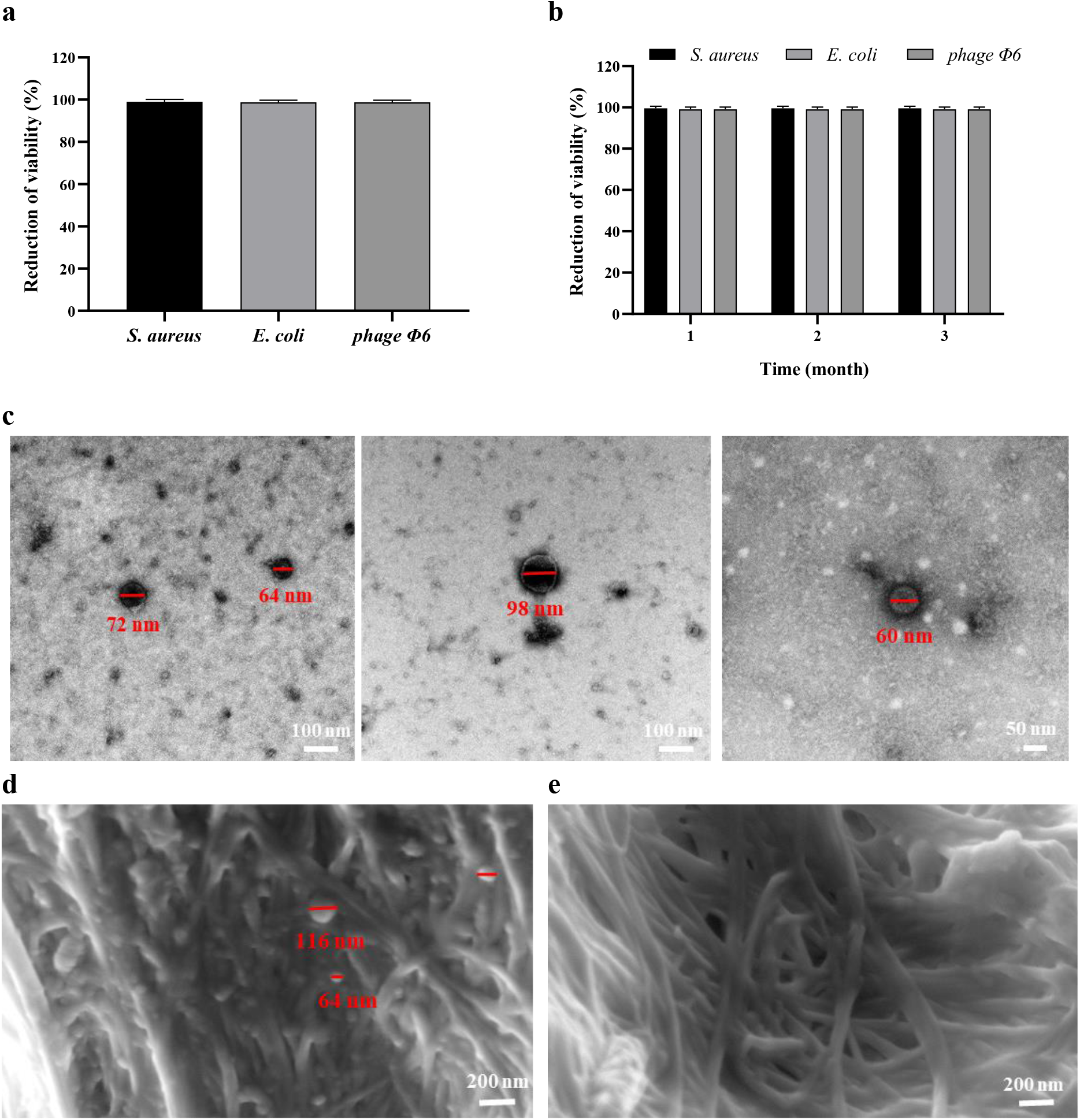
Antibacterial and antiviral properties of LPP-Ar-BC-bM. Antibacterial and antiviral activity of LPP-Ar-BC-bM after **a)** < 1 h from functionalization, **b)** 1, 3 and 3 month of storage. SEM micrographs of **c)** phage Φ6; **d)** phage Φ6 attached to the surface of control BC-bM; **e)** surface of LPP-Ar-BC-bM. Data in **a)** and **b)** were presented as a mean ± standard error of the mean (SEM). SEM micrographs were obtained at 10000x and 60000x magnification. Phage Φ6 diameters were measured using ImageJ software and are marked in red.

Additionally, we also used SEM to check for the presence of Φ6 phages on the surface of LPP-Ar-BC-bM and BC-bM. First, we confirmed that we were able to visualize phage Φ6 particles with the expected diameter (Fig. 5c). However, while we were able to observe phage particles on the control BC-bM, we did not note the presence of phages on the surface of LPP-Ar-BC-bM, which is consistent with the strong antiviral effect observed (Fig. 5d,e).

### Screening for potential cytotoxicity of LPP-Ar-BC-bM

LPP-Ar treatment resulted in marked antibacterial and antiviral effects of LPP-Ar-BC-bM. As a result, we wanted to screen for any potential cytotoxicity of the functionalized material against mammalian cells. Thus, LPP-Ar-BC-bM and BC-bM (used as a control) were evaluated using extract and direct contact assays, based on ISO 10993-5. In the extract assay, there was no evidence of cytotoxicity of any of the tested samples after 24 h of the culture of L929 murine fibroblasts with extracts (Fig. 6a). There was no difference in viability between the LPP-Ar-BC-bM and control BC-bM. The morphology of L929 cells was not altered (Fig. 6c).

**Fig. 6.**
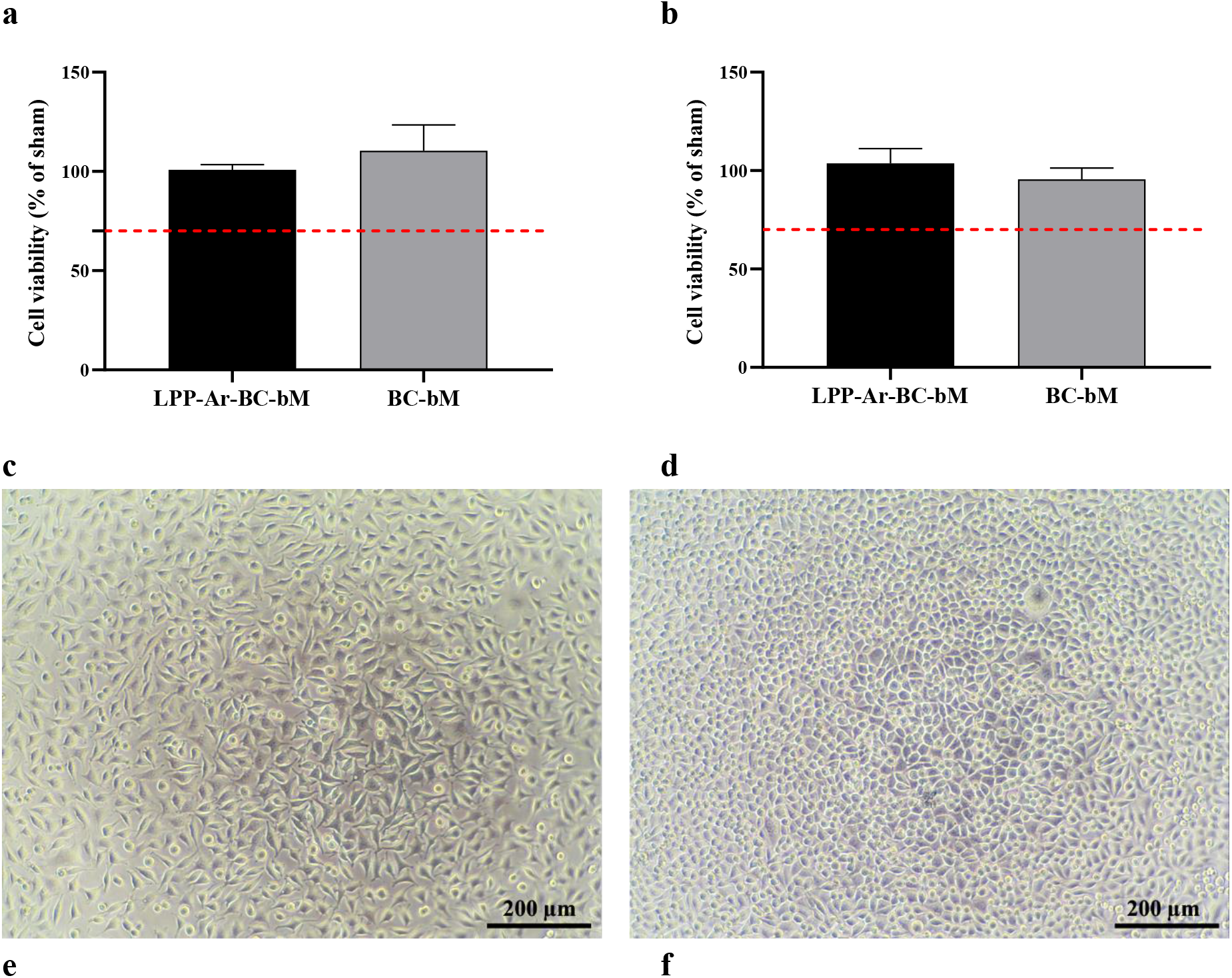
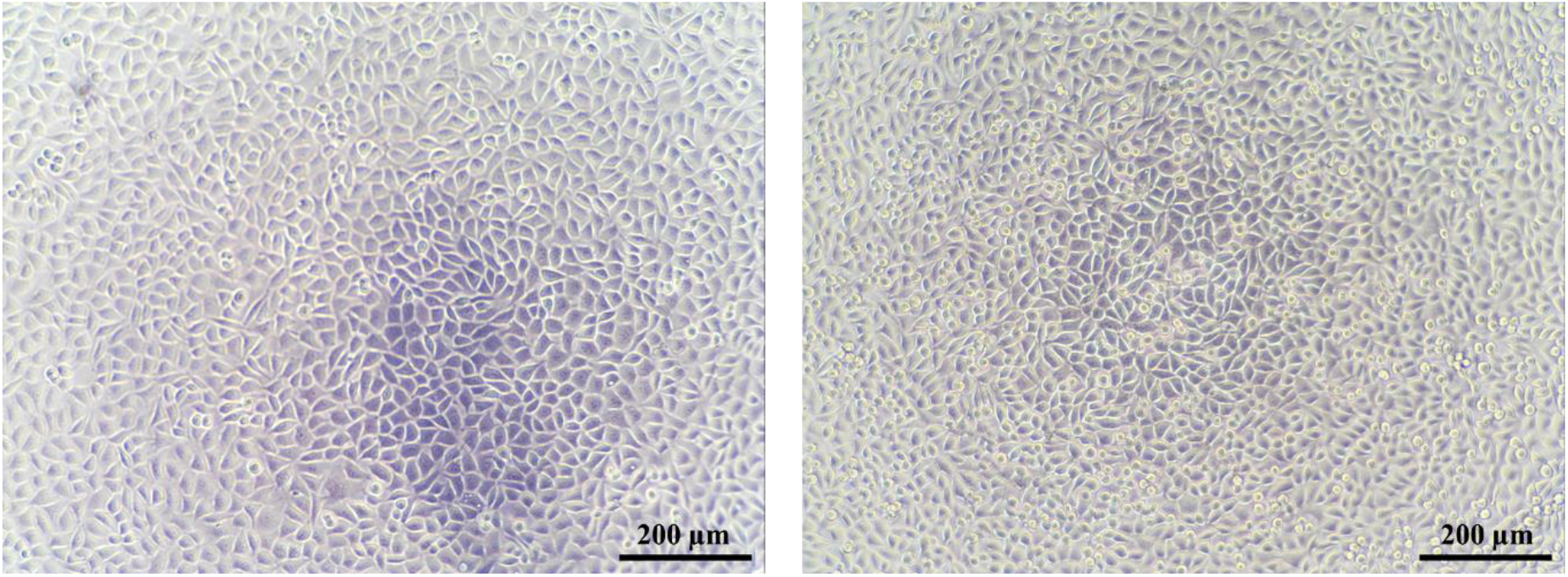
Screening for potential cytotoxicity of LPP-Ar-BC-bM. Viability of L929 fibroblasts after 24 h of culture **a)** with extract of LPP-Ar-BC-bM and BC-bM; **b)** in direct contact with discs of LPP-Ar-BC-bM and BC-bM for 24 h. Representative micrographs of L929 cells: **c)** 24 h after seeding (10000 cells per well); **d)** after 24 h of incubation with sham extract; **e,f)** after 24 h incubation with extracts of LPP-Ar-BC-bM and BC-bM, respectively. Data in **a)** and **b)** were presented as mean ± standard error of the mean (SEM). Red line indicates the cytotoxicity threshold from ISO 10993-5. There were no statistically significant differences between the materials (p<0.05).

Likewise, the direct contact assay, where discs of samples were placed directly on top of L929 fibroblast cells for 24 h, also did not show any difference between LPP-Ar-BC-bM and control BC-bM (Fig. 6b). We did not see altered morphology or reduced cell numbers when imaging wells without removing discs for the resazurin viability assay. Both directly under the discs and at the edges of the discs, cells were normal and dense (Fig. 6d).

### Evaluation of adsorption capacity of LPP-Ar-BC-bM

While our goal was to use LPP-Ar treatment to obtain the materials with antimicrobial and antiviral activity, we wanted to ensure that LPP-Ar treatment, which affected microstructure and porosity, did not reduce the ability of the materials to trap particles via adsorption. In fact, the results showed a modest increase in bacterial adsorption capacity for LPP-Ar-BC-bM, as compared to the control BC-bM (Table 2). Meanwhile, for the case of our model viral pathogen, phage Φ6, the LPP-Ar-BC-bM exhibited nearly 2-fold greater adsorption capacity, as compared to the control BC-bM.

**Table 2.**
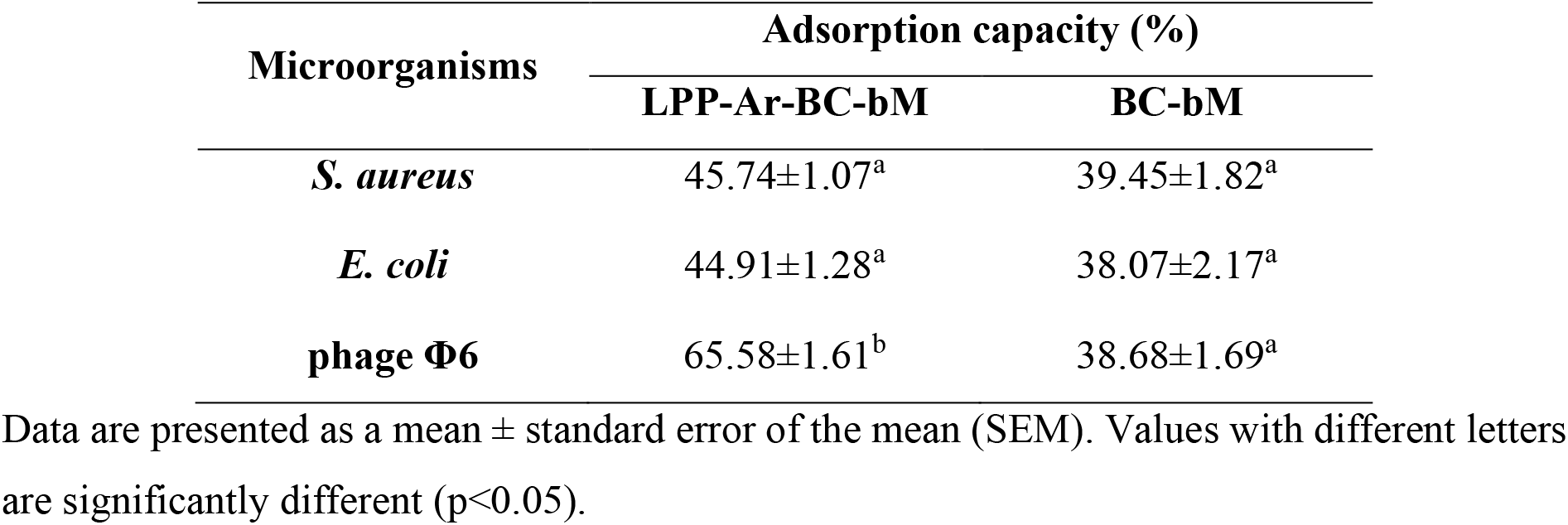
Adsorption capacity of LPP-Ar-BC-bM vs. BC-bM.

| Microorganisms | Adsorption capacity (%) |  |
| --- | --- | --- |
|  | LPP-Ar-BC-bM | BC-bM |
| *S. aureus* | 45.74±1.07<sup>a</sup> | 39.45±1.82<sup>a</sup> |
| *E. coli* | 44.91±1.28<sup>a</sup> | 38.07±2.17<sup>a</sup> |
| phage $\Phi 6$ | 65.58±1.61<sup>b</sup> | 38.68±1.69<sup>a</sup> |

Data are presented as a mean ± standard error of the mean (SEM). Values with different letters are significantly different ( $p < 0.05$ ).

### Evaluation of filtration efficiency and airflow resistance of LPP-Ar-BC-bM arranged in layers

Having shown that the obtained LPP-Ar-BC-bMs possess antimicrobial and antiviral properties, we assessed bacterial filtration efficiency (BFE) and viral filtration efficiency (VFE) in accordance with EN 14683+AC: 2019-09. Our results indicated that 1 layer of LPP-Ar-BC-bM provides on average ~80% bacterial filtration efficiency (BFE), but by arranging them in three layers, it was possible to ensure > 99% of BFE (Table 3). Importantly, for the case of phage Φ6, intended to model respiratory pathogens like SARS-CoV-2, just one layer of LPP-Ar-BC-bM ensured VFE above 99%.

**Table 3.**
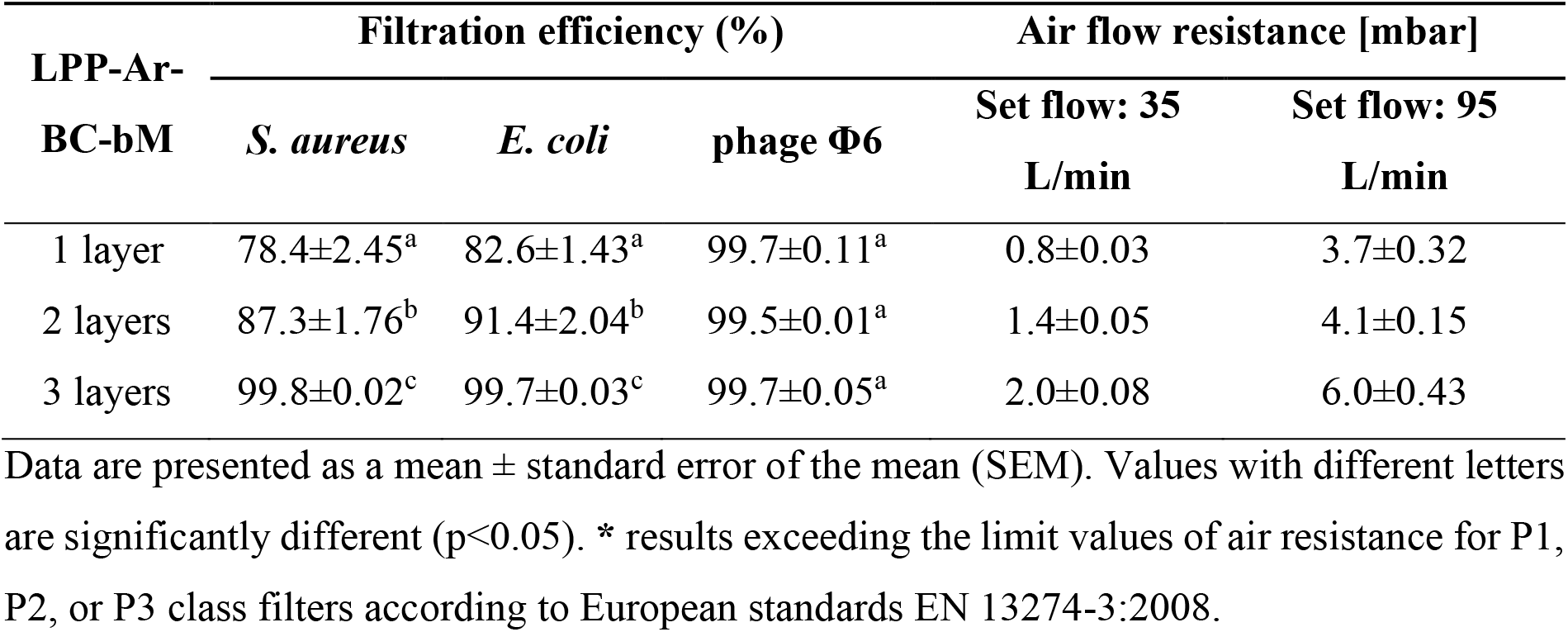
Filtration efficiency and airflow resistance of LPP-Ar-BC-bM.

Importantly, it was also confirmed that two or three layers of LPP-Ar-BC-bM still met the airflow resistance requirements for P1, P2, or P3 class filters according to European standards EN 13274-3:2008 *Respiratory Protective Devices - Methods of Test - Part 3: Determination of Breathing Resistance* (Table 3).

### User experience testing of prototype masks with LPP-Ar-BC-bM filter

As a final proof-of-concept, to demonstrate the potential of the developed functionalized materials to be used in PPE, we prepared 80 NanoBioCell masks and had 80 volunteer medical professionals wear them for 3 h. The participants then completed a user experience survey. In terms of comfort and fit, over all categories, only ~1% of participants rated the NanoBioCell masks as “poor”, with the vast majority (~60%) scoring the masks as “good” (Fig. 7a, Supplementary Fig. 6). Encouragingly, more participants (~25%) rated the prototype masks as “very good” as compared to “average” (~14%). In terms of moisture absorption, 81% of respondents described the NanoBioCell as “dry” after 3 h of use and only ~1% rated it as “wet” (Fig. 7b).

**Fig. 7.**
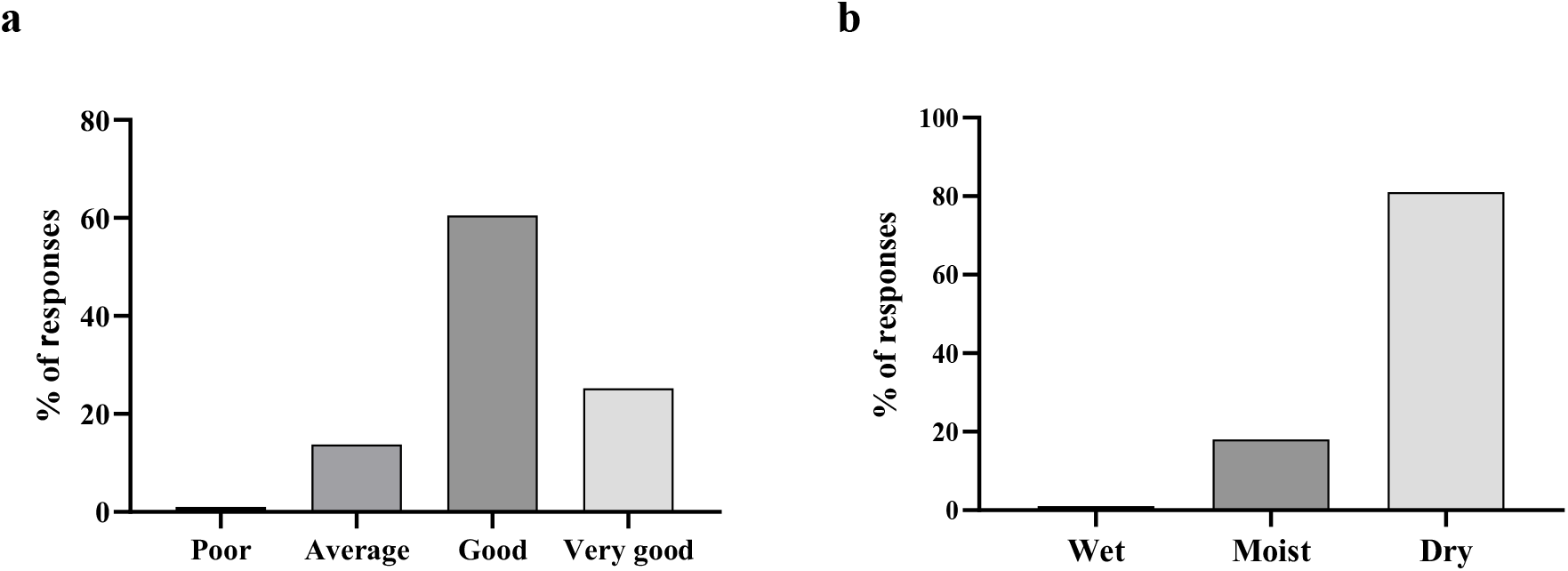
Results of user experience survey after 3 h of wearing the NanoBioCell mask. **a)** averaged comfort scores while donning and removing, making head movements, speaking, and breathing; **b)** moisture absorption after 3 h of using the NanoBioCell mask. The results were presented as % of responses against the number of respondents (n=80).

## Discussion

We hypothesized that the BC is well suited to function as a biobased, biodegradable filter material thanks to its unique nanoscale fibril morphology. Further, we aimed to use the LPP-Ar to improve antimicrobial and antiviral properties, without the need for harsh solvents or generating other waste. Ultimately, our goal was to develop a strategy to obtain biobased, biodegradable filters with excellent filtration efficiency parameters and antibacterial and antiviral properties that could be used in masks and other PPE. In this regard, such filters could act as sustainable, indispensable protective measures during the current SARS-CoV-2 pandemic or any future respiratory pathogen pandemic.

Despite the porous nanostructure of BC, its dense network makes obtaining sufficiently low airflow resistance particularly challenging^16^. In fact, we were not able to obtain sufficiently low airflow resistance values from dried BC pellicles to make them suitable as filters. Therefore, we decided to explore a homogenization and lyophilization process to obtain BC-bMs. The goal was to reduce the density and thus reduce airflow resistance. We tested different process conditions (BC content in the homogenized pulp and the pulp volume to surface ratio) that yielded materials with different densities and thicknesses. The BC-bMs were then tested for their airflow resistance in accordance with the European standard EN 13274-3:2008 for testing respiratory PPE. As a result, we were able to optimize process parameters to obtain excellent airflow resistance results while maintaining the homogeneous, consistent, three-dimensional structure of the BC-bMs.

Next, we aimed to obtain the antibacterial and antiviral properties of our BC-bMs using LPP-Ar. In contrast to various “wet” textile processing methods, LPP treatment is a solvent-free (“dry”) process that generates no waste, making it environmentally friendly and “green”^13^. However, the scope of research into LPP-based modification of BC has thus far been limited to enhancing cell affinity, adhesion, or change wettability^37–41^.

To guide the optimization of LPP-Ar process conditions, we used ATR-FTIR spectroscopy. With increasing time of LPP-Ar functionalization, LPP-Ar-BC-bMs showed an increasing absorbance band at 1720 cm^−1^ assigned to the carbonyl groups of the carboxyl group. The presence of these groups is related to the oxidation of primary alcohols, mainly C6-OH, presented in the structure of anhydrous glucose units^42,43^. The increase in absorbance at 1720 cm^−1^ plateaued after 10 min of LPP-Ar treatment, while 30 min of treatment resulted in visible yellowing, suggesting some thermal effects. Because it has been established that reactive species, radicals, charged particles and/or other atomic excited species generated during LPP-Ar could affect the crystallinity and thermal stability of the BC^44,45^, we used 2D ATR-FTIR spectroscopy, XRD, and TGA to carefully assess the crystallinity and thermal properties of the obtained materials. Collectively, the results indicated that after 30 min of LPP-Ar treatment the crystallinity and thermal stability of LPP-Ar-BC-bMs were decreased. Therefore, we concluded that 10 min of LPP-Ar treatment was optimal for BC-bMs functionalization. Importantly, we used ATR-FTIR spectroscopy to confirm that the LPP-Ar induced increase in absorbance at 1720 cm^−1^ was not lost over 1, 2, or 3 months of storage at room temperature in a desiccator.

Afterward, SEM imaging of LPP-Ar-BC-bM showed that the plasma process resulted in greater, although more heterogenous, porosity. This is consistent with previous reports by Gasi et al. that APP treatment induced the formation of valleys in the surface of polyamide fabric with depth and size associated with the interaction with the plasma^46^. Similarly, using SEM, Puppolo et al. showed that treating PES membranes with dichloromethane plasma resulted in surface restructuring and an increase in roughness, porosity, and rigidity^47^. Finally, Kutová et al. observed higher roughness and porosity of BC modified with argon plasma that was correlated with improved adhesion of keratinocytes^37^.

Having optimized the LPP-Ar process from a physicochemical point of view, we set out to assess whether the process resulted in improved antibacterial and antiviral properties. We confirmed that after 10 min of LPP-Ar treatment, LPP-Ar-BC-bM displayed strong antimicrobial activity against Gram-positive *S. aureus* and Gram-negative *E. coli* bacteria, as well as strong antiviral activity against bacteriophage Φ6 - a non-pathogenic model for SARS-CoV-2. Our results are consistent with earlier reports that LPP-Ar improves the antibacterial activity of different types of polymers^27,34,35,48^. Although low-temperature plasma (LTP) is a method commonly used for polymer surface modification, the mechanism responsible for antimicrobial activity (especially in the case of porous surfaces) remains to be fully elucidated^27^. LTP treatment may change oxidation, nitration, hydrolyzation, and/or amination of the surface, which can affect both prokaryotic and eukaryotic cell attachment and viability^27,49^.

In contrast to the well-established antimicrobial effect of LPP treatment, there has been limited research into antiviral activity. A direct antiviral effect of LPP-O_2_ 100% and LPP-Ar 80% + O_2_ 20% has been demonstrated towards bovine viral diarrhea virus and the porcine parvovirus, which are surrogates of human hepatitis C virus and human parvovirus B19, respectively^50,51^, but this makes it a sterilization process, rather than a lasting functionalization. However, in this work we showed that 10 min of LPP-Ar could yield LPP-Ar-BC-bM with strong antiviral activity: we observed a >99% reduction in the activity of phage Φ6, as compared to BC-bM. Further, the effect was not diminished over 3 months of storage. Phage Φ6 was chosen as a model virus, because it is of comparable size and has a somewhat similar lipid envelope to SARS-CoV-2, which makes it a suitable non-pathogenic surrogate for studying the surface and air survival of pathogenic viruses, including SARS-CoV-2^36^. Using SEM imaging we did not observe phage Φ6 particles on the surface of LPP-Ar-BC-bM, in contrast to the BC-bM. This suggests that direct contact with the surface of LPP-Ar-BC-bM may cause the lipid envelope of phage Φ6 to disintegrate, resulting in phage inactivation^36^.

Taking into consideration the strong antibacterial and antiviral properties of LPP-Ar-BC-bM, we screened for potential cytotoxicity against eukaryotic, mammalian cells using L929 murine fibroblasts. The results of both extract and direct contact assays based on ISO 10993-5 showed no reduction in L929 viability reduction, no changes in cell morphology, and no differences between LPP-Ar-BC-bM and BC-bM. Therefore, we conclude that LPP-Ar treatment did not have any adverse effect on the cytocompatibility of the materials and that the obtained LPP-Ar-BC-bM can be considered non-toxic.

In order to demonstrate that the developed LPP-Ar-BC-bM is suitable for respiratory PPE, such as surgical masks, we tested filtration efficiency in accordance with ISO 14683+AC:2019-09. This standard defines the degree of filtration efficiency required by distinct types of medical face masks and other medical filtration materials. For type I medical masks, the bacterial filtration efficiency (BFE) must be ≥ 95%, while for type II and IIR it must be ≥98%. Our results showed that by using three layers of LPP-Ar-BC-bM, we could obtain a BFE >99% for *S. aureus* and *E. coli*. Further, and perhaps more importantly in the context of the present SARS-CoV-2 pandemic, the viral filtration efficiency (VFE) was >99% with single layer LPP-Ar-BC-bM. It is important to note that three layers of LPP-Ar-BC-bM still met the requirements of the EN 13274-3: 2008 standard for airflow resistance (6.0±0.43 mbar resistance at an airflow of 95 L/min). According to EN 13274-3:2008 standard, at an airflow of 95 L/min, the air resistance of respiratory PPE should not exceed 6.1 mbar for a class P1 filter, 6.4 mbar for a class P2 filter (equivalent to a N95 respirator), and 8.2 mbar for class P3 filter.

Overall, we conclude that the high filtration efficiency of our three-layers LPP-Ar-BC-bM filter is the result of the combination of the unique structure of BC-bM and LPP-Ar induced changes in isoelectric charge, polarity, reactivity, and wettability, which increase biomolecule adsorption^52–54^. It has been previously shown that LTP-modified surfaces have an increased ability to adsorb proteins present in mammal and bacterial cells^55–57^. In fact, Griffin et al. confirmed that LPP-Ar surface modification correlated with the higher adsorption of such proteins, as compared to LPP treatment with oxygen or nitrogen^57^. For the case of enveloped viruses like SARS-CoV-2 (and the surrogate we used, phage Φ6) the virus is protected by an outer lipid bilayer with a high density of proteins, such as the crucial spike protein^58–60^. We hypothesize that it is interactions with these proteins that may explain the high ability of LPP-Ar-BC-bM filters to both adsorb and deactivate viral particles.

As a final proof-of-concept that our filters could be used in PPE filtration devices, we prepared 80 NanoBioCell masks and asked volunteer medical staff to wear them for 3 h. The majority of scores were above the score “average”, which indicates that, from a comfort perspective, the NanoBioCell masks were comparable to, or better than, the existing PPE used by the participants. Only ~1% of responders rated the masks as “poor”, which may have been the result of imperfect internal quality control, because members of the research team and not a professional company made not just the filters but the entire masks. When combined with the fact that the LPP-Ar-BC-bM filters had excellent BFE and VFE and are fully biobased and biodegradable, these results establish the excellent potential of the developed process and materials to be used for PPE products.

In summary, in the current work, we showed that it is possible to fabricate air filters using bacterial cellulose (BC) by using homogenization and lyophilization. Further, the filters could be functionalized using low-pressure argon plasma (LPP-Ar), which markedly improved antimicrobial and antiviral properties. Such three-layers LPP-Ar-BC-bM filters provided strong antibacterial and antiviral properties, as well as bacterial and viral filtration efficiencies >99%. At the same time, the three-layers LPP-Ar-BC-bM filters met the airflow resistance requirements of the EN 13274-3: 2008 standard for respiratory PPE. Further, despite the potent antimicrobial and antiviral effect, we did not observe any indication of cytotoxicity caused by the LPP-Ar-BC-bM. Importantly, by combining the biotechnological BC production with “green” low-temperature plasma functionalization, the entire process is environmentally friendly. Our filters do not require the use of any harmful solvents and produce no hazardous waste. Finally, as a proof-of-concept, we were able to prepare 80 NanoBioCell masks and ~85% of volunteer medical staff assessed them as “good” or “very good” in terms of comfort. As a result, we conclude that our LPP-Ar-BC-bM filters can be used in PPE face masks and respirators, offering a biobased, biodegradable, sustainable and safe alternative to existing materials. Further, we believe that with scale-up, the developed technology could be used for large indoor air filtration systems, such as those in hospitals or schools.

## Materials and methods

### Preparation of BC-based material

In the first stage, BC was biosynthesized using a reference strain of *Komagataeibacter xylinus* (American Type Culture Collection – ATCC 53524). For this purpose, bacterial suspension of 3×10^5^ colony forming units per milliliter (CFU/mL) was used to inoculate 100 mL of Hestrin-Schramm (H-S) culture medium in Petri dishes (15 cm diameter) (Becton Dickinson and Company, USA). BC biosynthesis was then conducted for 7 days at 28 °C. The obtained BC pellicles were then purified to remove bacteria and culture medium components by using a 0.1 M sodium hydroxide solution at 80 °C for 90 min, followed by rinsing with distilled water until the pH was neutral^61^.

In the next stage, the purified BC pellicles were homogenized using a blender (Perfectmix + BL811D38, Tefal, France) with the addition of distilled water at a weight ratio of 1:1, 1:2, or 2:1. Once homogeneous BC pulp was obtained, it was poured into square Petri dishes (120 × 120 × 17 mm) at a volume of 60 mL, 80 mL, or 100 mL. The pulp was then frozen at −18 °C or −80 °C for 24 h and then lyophilized at −60 °C and 0.1 mBar (Alpha 1-2 Ldplus, Christ, Germany) to yield BC-based materials (referred to as “BC-bMs”) (Fig. 8).

**Fig. 8.**
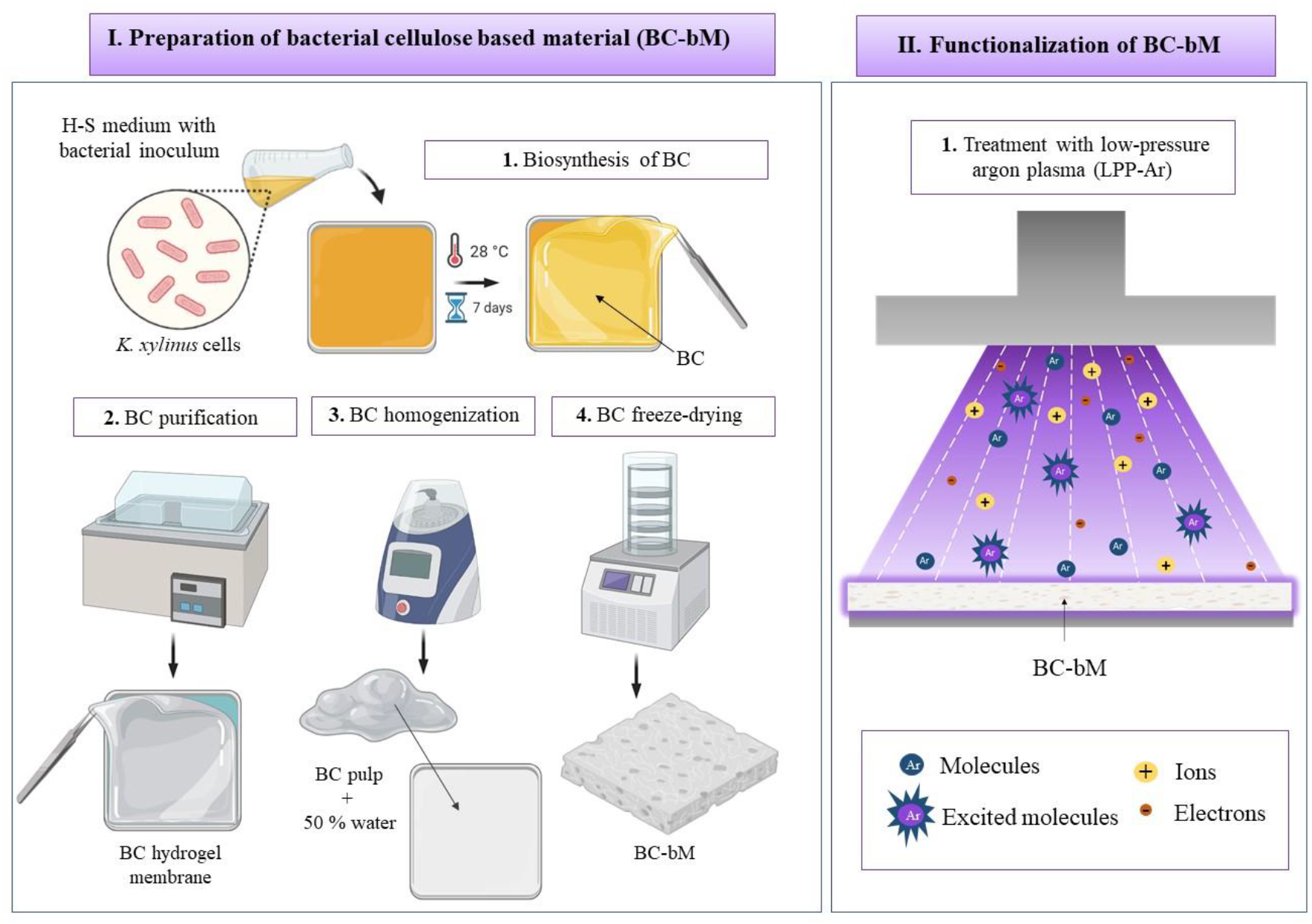
Scheme of preparation and functionalization of BC-bMs.

### Evaluation of BC-bMs

#### Macrostructure analyses

The macro-morphological structure of the surfaces, as well as cross-sections of the BC-bMs were examined using a stereoscopic microscope (Leica S9i, Leica Microsystems, Wetzlar, Germany).

#### Airflow resistance analyses

Airflow resistance was assessed according to the European standard EN 13274-3:2008 *Respiratory Protective Devices – Methods of Test – Part 3: Determination of Breathing Resistance*, which specifies the general procedure for measuring the breathing resistance of BC-bMs for respiratory protective devices (Supplementary Fig. 7). The tests were conducted at airflow rates of 30 L/min and 95 L/min, corresponding to a respiratory minute volume during moderate and strenuous activity.

### Functionalization of BC-bM

The BC-bMs were functionalized with a low-pressure argon plasma (LPP-Ar) using HPT-100 Benchtop Plasma Treater (Henniker Plasma, UK). The gas flow and power were held constant at 10 sccm (chamber pressure ~0.6 mbar) and 100% power (100 W). These process parameters were selected in cooperation with a technical support staff of Henniker Plasma and yielded stable, well-distributed plasmas within the chamber. The treatment time was varied from 1 to 30 min. After LPP-Ar treatment, the functionalized BC-bMs (referred to as “LPP-Ar-BC-bMs”) were stored at room temperature in a desiccator until further analysis.

### Evaluation of LPP-Ar-BC-bM

#### Attenuated Total Reflectance Fourier Transform Infrared Spectroscopy analysis

Attenuated Total Reflectance Fourier Transform Infrared Spectroscopy (ATR-FTIR) was used to characterize functional groups and detect changes in chemical composition in LPP-Ar-BC-bMs. The measurements were performed using ALPHA FT-IR Spectrometer (Bruker Co., Germany) with a DTGS detector and the platinum-ATR-sampling module with a robust diamond crystal and variable angle incidence beam. For each sample, 32 scans at 2 cm^−1^ resolution were recorded over the spectral range of 4000 - 400 cm^−1^. The spectra were processed by baseline correction, smoothed with a polynomial Savitzky-Golay filter, and normalized to band area at 1161 cm^−1^ using the SpectraGryph 1.2 software package. Processed spectra were then analyzed using two-dimensional correlation spectroscopy in OriginPro2021 software. The BC-bM was used as a control. Additionally, ATR-FTIR was used to assess the stability of functionalization after 1, 2, and 3 months of storage of LPP-Ar-BC-bMs at room temperature in a desiccator.

#### X-ray diffraction analysis

X-ray diffraction analysis (XRD) using a D5005 X-ray diffractometer (Bruker Siemens, USA) was used to assess the crystallinity of LPP-Ar-BC-bMs. A diffraction angle 2 θ was measured from 5° to 70° with a step size of 0.04° using Cu-kα radiation at 40 kV and 40 mA. Crystallinity (%) was calculated by dividing the area of the crystalline peaks by the total area under the curve from 2θ 5° to 30° ^**62**^, using the Diffrac.eva software by Bruker (USA). The BC-bM was used as a control.

#### Thermogravimetric analysis

The thermal stability of LPP-Ar-BC-bMs was evaluated using a Perkin Elmer TGA 8000 thermogravimetric analysis (TGA) system. Samples were conditioned at room temperature before measurements without a pre-heating cycle, to obtain water evaporation data. Then, during the experiment, samples (~6 mg) were placed in ceramic sample pans and were heated from 30 °C to 700 °C, at a heating rate of 10 °C/min under nitrogen atmosphere. The BC-bM was used as a control.

### Evaluation of LPP-Ar-BC-bM functionalized under optimal conditions (10 min LPP-Ar)

#### Scanning electron microscopy analysis

To examine the microstructure of LPP-Ar-BC-bM using scanning electron microscopy (SEM), samples were first sputtered with Au/Pd (60:40) using Leica EM ACE600 sputter (Leica Microsystems, Wetzlar, Germany). The surfaces of the sputtered samples were then examined using Auriga 60 SEM (Zeiss, Oberkochen, Germany). Pore diameters were determined using ImageJ software (NIH) software integrated with the Auriga 60 SEM. The BC-bM was used as a control.

#### Analysis of antimicrobial activity

The antimicrobial activity of LPP-Ar-BC-bM was tested against Gram-positive *Staphylococcus aureus* ATCC 6538 and Gram-negative *Escherichia coli* ATCC 25922 in accordance with ISO 20743:2013 *Textiles – Determination of antibacterial activity of textile products*. Briefly, 0.4 g of LPP-Ar-BC-bM was transferred to a Petri dish. Next, 200 μL of bacterial suspension (3×10^5^ CFU/mL) in Brain Heart Infusion (BHI, Graso, Poland) broth medium was applied to each tested sample and incubated for 24 h at 37 °C. After incubation, the samples were placed in 50 mL plastic tubes (Polypropylene Conical Centrifuge Tube, Becton Dickinson and Company, USA) with 20 mL of phosphate-buffered saline (PBS, Sigma Aldrich, Germany) and vortexed for 60 s to remove the bacterial cells attached to the sample surfaces. The number of workable bacterial cells in the suspension was determined by quantitative plating on BHI agar medium after 24 h of incubation at 37 °C. The results are presented as a reduction of viability (%) compared to BC-bM used as a control. Additionally, antibacterial tests were performed after 1, 2, and 3 months of storage of LPP-Ar-BC-bMs at room temperature in a desiccator to assess the stability of functionalization.

#### Analysis of the antiviral activity

To assess potential antiviral activity of LPP-Ar-BC-bM, we used *Pseudomonas* phage Φ6 (German Collection of Microorganisms and Cell Cultures GmbH (Deutsche Sammlung von Mikroorganismen und Zellkulturen DSM 21518). The phage Φ6 is an enveloped bacteriophage, non-pathogenic to humans, that can be used as a model surrogate for studying the surface and air survival of pathogenic viruses, including SARS-CoV-2^36^.

Testing was conducted following ISO 18184:2019 *Determination of the activity of antiviral textiles – Absorption method*. Briefly, 0.4 g of LPP-Ar-BC-bM was placed in a Petri dish and 200 μL of phage lysate at concertation of 3×10^5^ plaque-forming units per milliliter (PFU/mL) in lysogeny broth medium (LB, Graso, Poland) was applied on each tested sample, followed by incubation for 24 h at 28 °C. After incubation, the samples were transferred into 50 mL conical tubes with 20 mL of SM buffer (50 mM Tris HCl, 100 mM NaCl, 8 mM MgSO4, pH 7.5, Merck, Germany) and vortexed for 60 s to remove phage Φ6 particles attached to sample surfaces. The number of active phage particles in the suspension was determined by plaque assay on LB agar medium after 24 h of incubation at 28 °C, with *Pseudomonas syringae* DSM 21482 used as the host. The results are presented as a mean reduction of viability (%) compared to the BC-bM used as a control. Additionally, samples were imaged using SEM as described previously to visualize phages Φ6 particles on and within the LPP-Ar-BC-bM as well as the BC-bM (used as a control) structure. Likewise, to confirm the long-term stability of the LPP-Ar functionalization, the antiviral activity of the LPP-Ar-BC-bM was tested also after 1, 2, and 3 months of storage.

#### Cytotoxicity screening

Extract assay. To screen for any potential cytotoxicity of LPP-Ar-BC-bM, an extract assay was performed following ISO 10993-5 *Biological evaluation of medical devices – Part 5: tests for in vitro cytotoxicity*. To obtain extracts, 3 discs (10 mm diameter, ~3 cm^2^) each of LPP-Ar-BC-bM and BC-bM (used as a control) were placed in wells of a 12-well plate and covered with 1 mL of growth medium: DMEM with 10% fetal bovine serum (FBS), 2 mM l-glutamine, 100 U/mL penicillin, and 100 μg/mL streptomycin (all cell culture reagents were purchased from Sigma-Aldrich, Poznań, Poland). As a sham control, 1 mL of media was pipetted into an empty well. Nitrile glove (Mercator Nitrylex Classic, Kraków, Poland) and polycaprolactone (PCL) (CAPA 6430, Perstorp, Sweden) were used as positive (toxic) and negative (non-toxic) assay controls, respectively (3 cm^2^ of material per mL of media)^63^. The plate was then incubated for 24 h at 37 °C in a cell culture CO_2_ incubator.

In parallel, a 96-well plate was seeded with 1×10^4^ L929 murine fibroblast cells per well and incubated for 24 h to allow for cell adhesion and spreading (L929 cell line was purchased from Sigma-Aldrich, Poznań, Poland and used between passages 10 and 30). Next, the media was aspirated and replaced with 100 μL of each extract, with 6 technical replicates performed per sample. The plate was then returned to the incubator and the cells were cultured for an additional 24 h, after which cells were examined using an inverted light microscope (Delta Optical IB-100, Poland). Cell viability was evaluated using a resazurin assay (25 μg/mL in complete growth media) (Riss et al., 2004), with fluorescence measurements performed using a fluorescent microplate reader (Synergy HTX, Biotek, USA) at 540 nm excitation and 590 nm emission. The results were expressed as a percent of cell viability relative to sham control and were calculated using the formula:

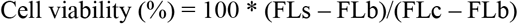

where FL is the fluorescence intensity (arbitrary units) and indexes s, b, and c refer to sample, blank, and sham control, respectively.

Direct contact assay. To further assess the cytocompatibility of LPP-Ar-BC-bM, we performed a direct contact assay based on ISO 10993-5 *Biological evaluation of medical devices – Part 5: tests for in vitro cytotoxicity* as in our previous work, with minor modifications^63^. L929 cells (passages 10–30) were seeded in wells of a 48-well plate at a density of 3×10^4^ cells per well and cultured for 24 h to allow for adhesion and spreading. For both LPP-Ar-BC-bM and BC-bM (used as a control), discs 8 mm in diameter (8 samples per material) were cut with a steel punch and soaked in complete growth media for ~5 min to ensure total saturation. For each sample, media was aspirated from the well with cells, and the disc was carefully placed directly on top of the cell monolayer, followed by the addition of 150 μL of complete growth media. As a sham control, media was aspirated and 300 μL of media was replaced. The plate was then returned to the cell culture incubator and cultured for 24 h. Cells were then imaged with discs in place using an inverted light microscope (Delta Optical IB-100, Poland). Next, 60 μL of resazurin stock (0.15 mg/mL in PBS) was added directly to each well and after 4 h of incubation, fluorescence measurements were performed, as described above, in “top read” mode, without removing the discs.

#### Evaluation of adsorption capacity

In order to evaluate the adsorption capacity, 0.4 g samples of LPP-Ar-BC-bM and BC-bM (used as a control) were placed in 50 mL plastic tubes (3.8 cm diameter) with either 20 mL of a bacterial suspension at a concentration of 3×10^5^ CFU/mL or phages at a concentration of 3×10^5^ PFU/mL. The prepared samples were then incubated for 1 h at 37 °C for the case of bacteria and 28 °C for the case of phage. After that time, the samples were rinsed with PBS (bacteria adsorption test) or SM buffer (phage adsorption test), to remove unattached bacteria or phage. Next, samples were transferred into new 50 mL plastic tubes containing 20 mL of PBS or SM buffer and vortexed for 60 s to remove bacterial cells or phage adsorbed on the surface of the samples. Finally, the number of adsorbed bacterial cells or phage particles in the suspension was determined as described previously in *2.5.2 Analysis of antimicrobial activity* and *2.5.3 Analysis of the antiviral activity*. The results are presented as a mean adsorption capacity (%) which was calculated using the following equation:

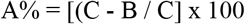

where:

A% – adsorption capacity of bacteria/phages;
C – initial number of bacterial cells/phage particles used for the test;
B – mean number of bacterial cells/phage particles released from test samples.

### Evaluation of filtration efficiency and airflow resistance of LPP-Ar-BC-bM arranged in layers

Next, to further develop our concept, we tested 1, 2, or 3 layers of LPP-Ar-BC-bM and assessed Bacterial Filtration Efficiency (BFE) and Viral Filtration Efficiency (VFE) following European standard *EN 14683 + AC: 2019-09 Medical masks – Requirements and test methods* (Supplementary Figure 8). The bacterial suspensions (3×10^5^ CFU/mL of *S. aureus* or *E. coli*) or phage lysate (3×10^5^ PFU/mL of phage Φ6) were aerosolized and delivered to the LPP-Ar-BC-bM samples (used as a filtration material) at a constant flow rate of 28.3 L/min. The filtration efficiency was calculated using the formula:

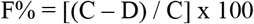

Where:

F% – filtration efficiency of bacteria/phages;
C – number of CFU or PFU in the control test (conducted without any filtration material);
D – number of CFU or PFU in the test with the filtration material (LPP-Ar-BC-bM).

At this stage of experiment, the airflow resistant of 1, 2 and 3 layers of LPP-Ar-BC-bM was assessed as described in *Airflow resistance analyses*.

### User experience testing of prototype masks consisting of LPP-Ar-BC-bM filter

As a final proof-of-concept, we prepared 80 prototype masks (referred to as “NanoBioCell”) consisting of 1 layer of LPP-Ar-BC-bM filter sandwiched between two layers of polylactic acid non-woven fabric (Gramix, Poland), (Supplementary Figure 9, Supplementary Figure 10). Masks were then tested by 80 volunteers consisting of the medical staff at two clinical hospitals in Szczecin, Poland, in accordance with institutional guidelines. For the testing, participants wore the prototype masks for 3 h and then completed a user experience survey prepared based on the European standard EN 13274-2 *Respiratory protective devices - Methods of test - Part 2: Practical performance tests*. The participants assessed the comfort and fit of the masks during donning, removal, and head movement, as well as the comfort of breathing, and comfort of speaking using a four-point scale: 1) poor, 2) average), 3) good, 4) very good. The participants also assessed the moisture absorbency of the prototype masks, rating the feeling of their masks after 3 h as dry, moist, or wet.

### Statistical analysis

Data are shown as means ± standard errors of the means (SEM) obtained from at least three different measurements (plus technical repetitions). Statistical differences between samples were determined by one-way analysis of variance (ANOVA) and Tukey’s post hoc test. All analyses were considered statistically significant when the *p*-value was less than 0.05. The statistical analyses were conducted using GraphPad Prism 9.0 (GraphPad Software Inc., USA).

## Data availability

Original datasets discussed in this publication have been deposited with link to Figshare database (https://figshare.com/articles/dataset/Argon_plasma-modified_bacterial_cellulose_filters_for_protection_against_respiratory_pathogens/19615236).

## Acknowledgements

This research was funded by the Regional Operational Program of the West Pomeranian Voivodeship, Grant No. Proto_lab/ K1/2020/U/11 and Proto_lab/K2/2021/U/7. We would like to thank Grace Law, Henniker Plasma, for technical advice and support regarding low pressure argon plasma parameters. We would also like to thank Xymena Stachurska for help in phage Φ6 lysate preparation.

## Author contributions

A.Ż., analyzed data, wrote the initial draft of the manuscript; D.C.J. M.Sz., R.D., P.S, A.J., S.G., methodology, investigation, data curation, and visualization; M.EF., funding acquisition, editing the manuscript; K.F., conceptually designed experiments, supervision, writing – review and editing the manuscript. All authors discussed the results and interpretations and reviewed the manuscript.

## Competing interests

The authors declare no competing interests.

## Notes

### Competing Interest Statement

The authors have declared no competing interest.

