## Supplementary data for "Argon plasma-modified bacterial cellulose filters for protection against respiratory pathogens"

**Results**

a b

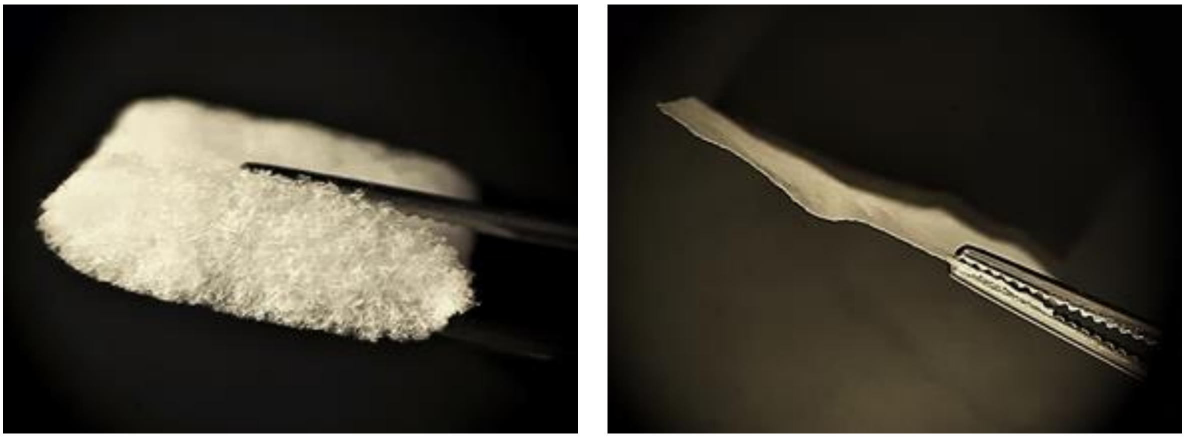

**Supplementary Figure 1.** Cross sections of **a)** BC-bM and **b)** dry BC pellicle.

Images were taken using stereoscopic microscope.

a b

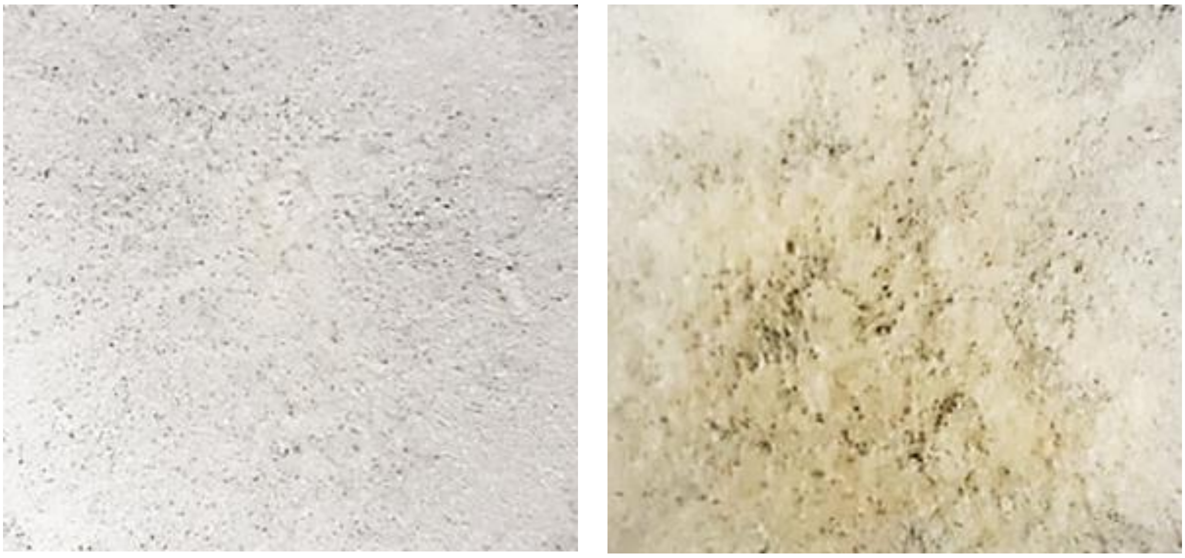

**Supplementary Figure 2.** The macro-morphological differences in BC-bMs depending on time of LPP-Ar functionalization process. **a)** 10 min; and **b)** 30 min.

Images were taken using stereoscopic microscope.

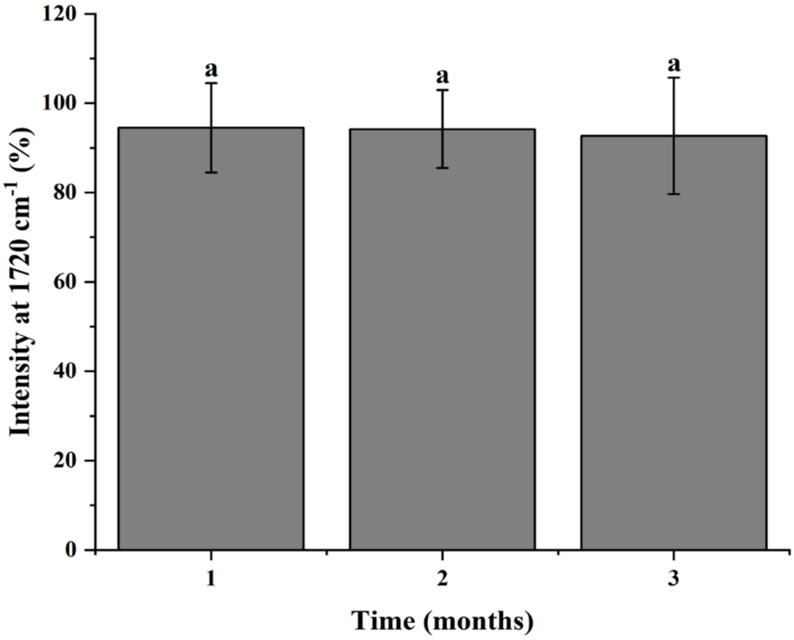


**Supplementary Figure 3.** The intensity of the ATR-FTIR absorbance band at 1720 $cm^{-1}$ over the course of 3 months of storage of LPP-Ar-BC-bMs.

The BC-bMs were treated for 10 min with LPP-Ar and then stored at room temperature in a desiccator. The intensity of the ATR-FTIR spectra absorbance band at 1720 $cm^{-1}$ was periodically checked. The intensity of the absorbance band in ATR-FTIR at 1720 $cm^{-1}$ was expressed as a percentage relative to the area of the band immediately after LPP-Ar treatment. The means with the same superscripts are not significantly different with a p-value above 0.05; error bars indicate standard error of the mean (SEM).

a b

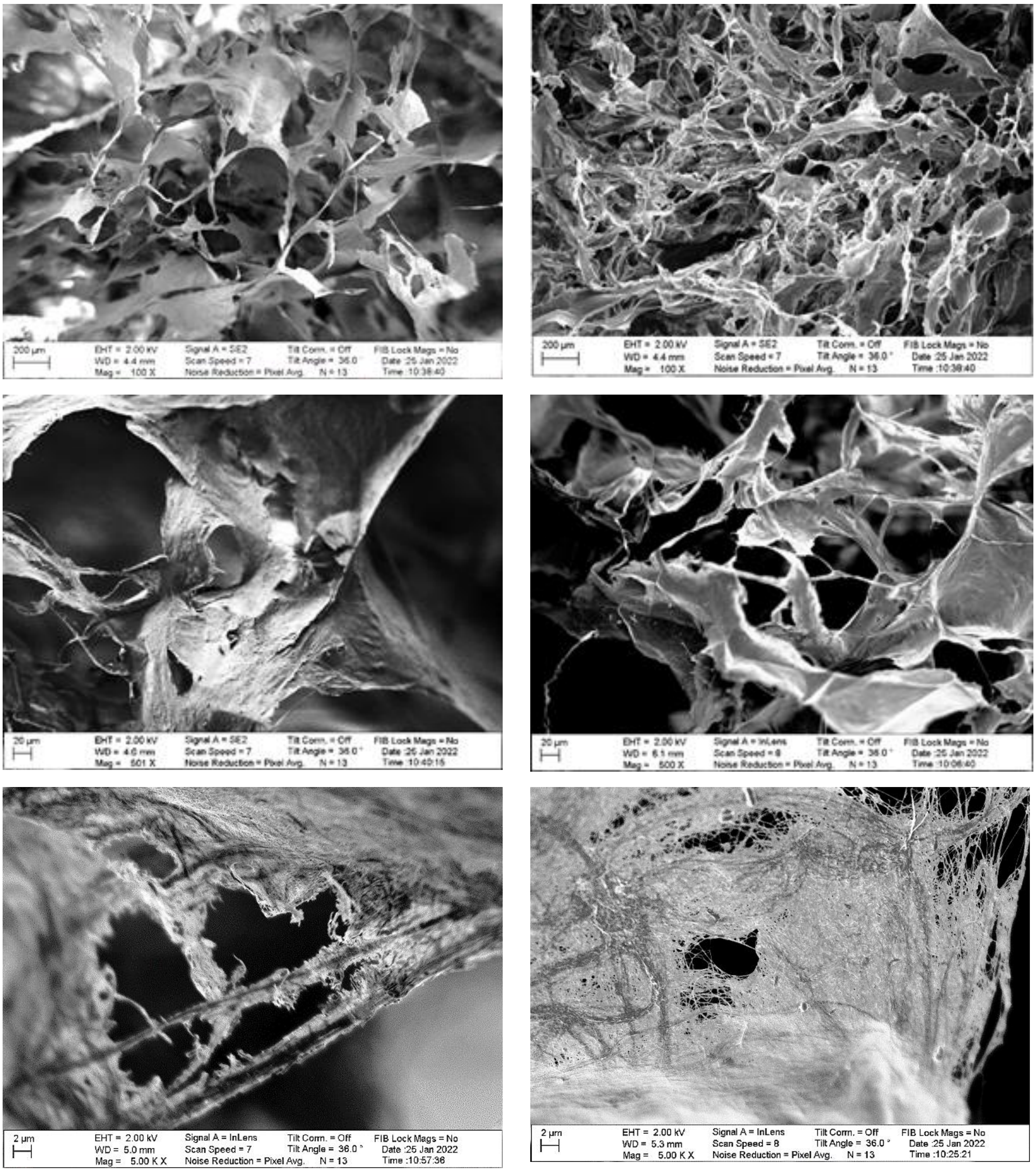

**Supplementary Figure 4.** Representative SEM micrographs of **a)** LPP-Ar-BC-bM and **b)** BC-bM.

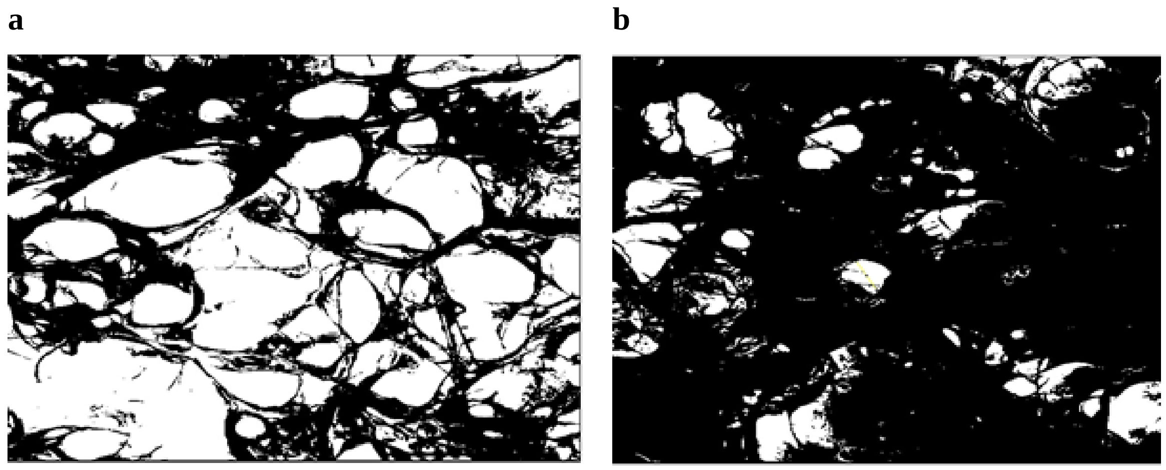


**Supplementary Figure 5.** Cellulose pores captured by ImageJ software. **a)** LPP-Ar-BC-bM; and **b)** BC-bM.

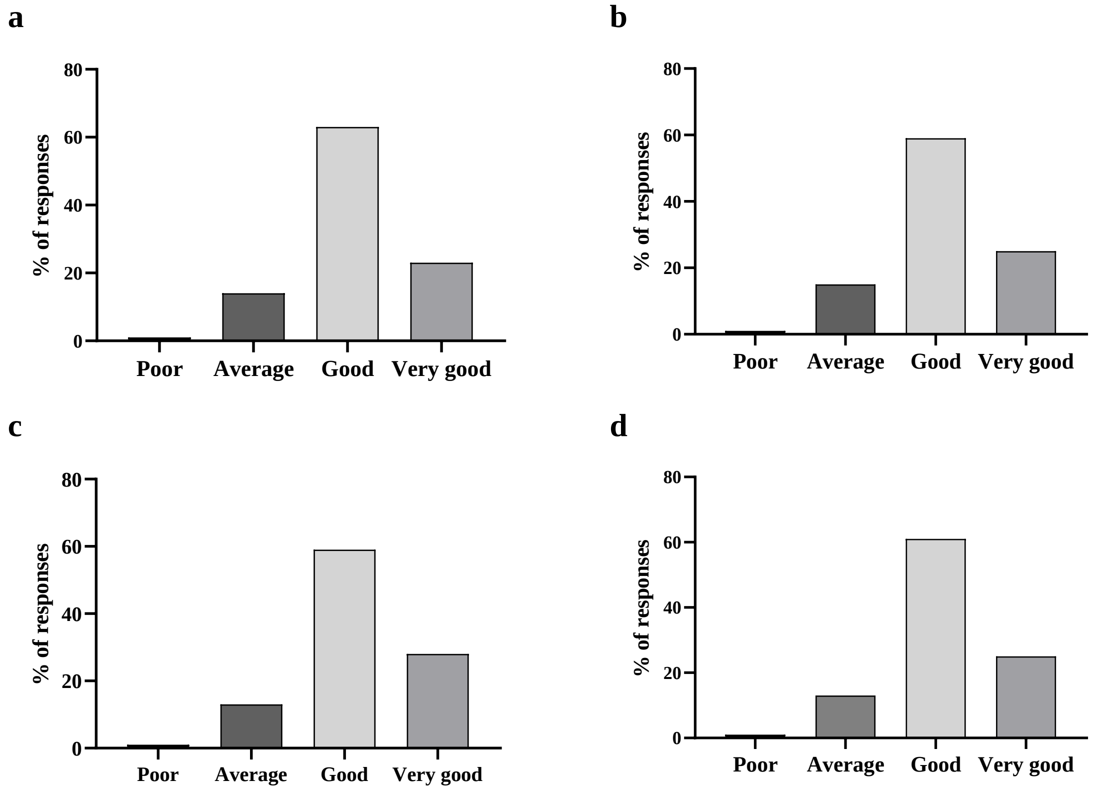


**Supplementary Figure 6.** Results of user experience survey after 3 h of wearing NanoBioCell mask. Comfort while: **a)** donning and removing; **b)** making head movements; **c)** speaking; **d)** breathing; and **e)** moisture absorption after 3 h of using NanoBioCell mask.

The results were presented as % of responses against the number of respondents (n=80).

**Materials and methods**

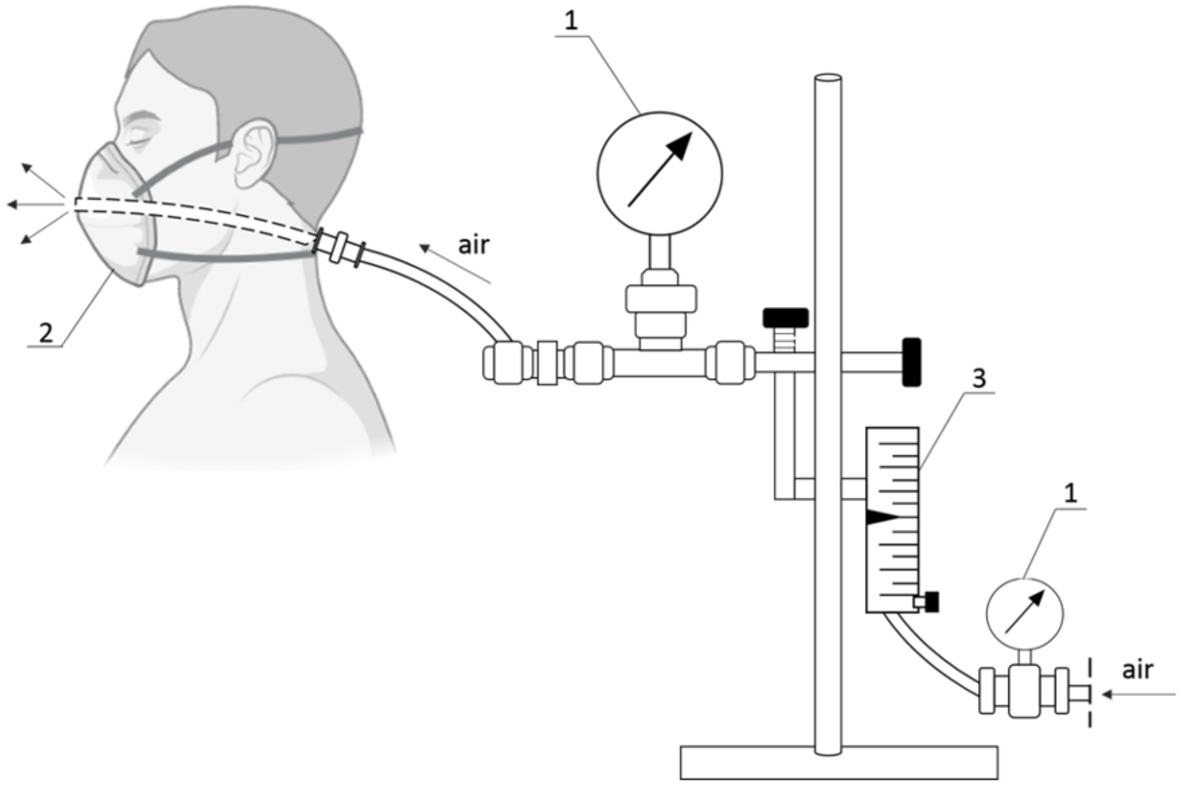


**Supplementary Figure 7**. Diagram of device used for measuring airflow resistance.

1 – manometer; 2 - mask housing for replaceable filter; 3 - airflow meter.

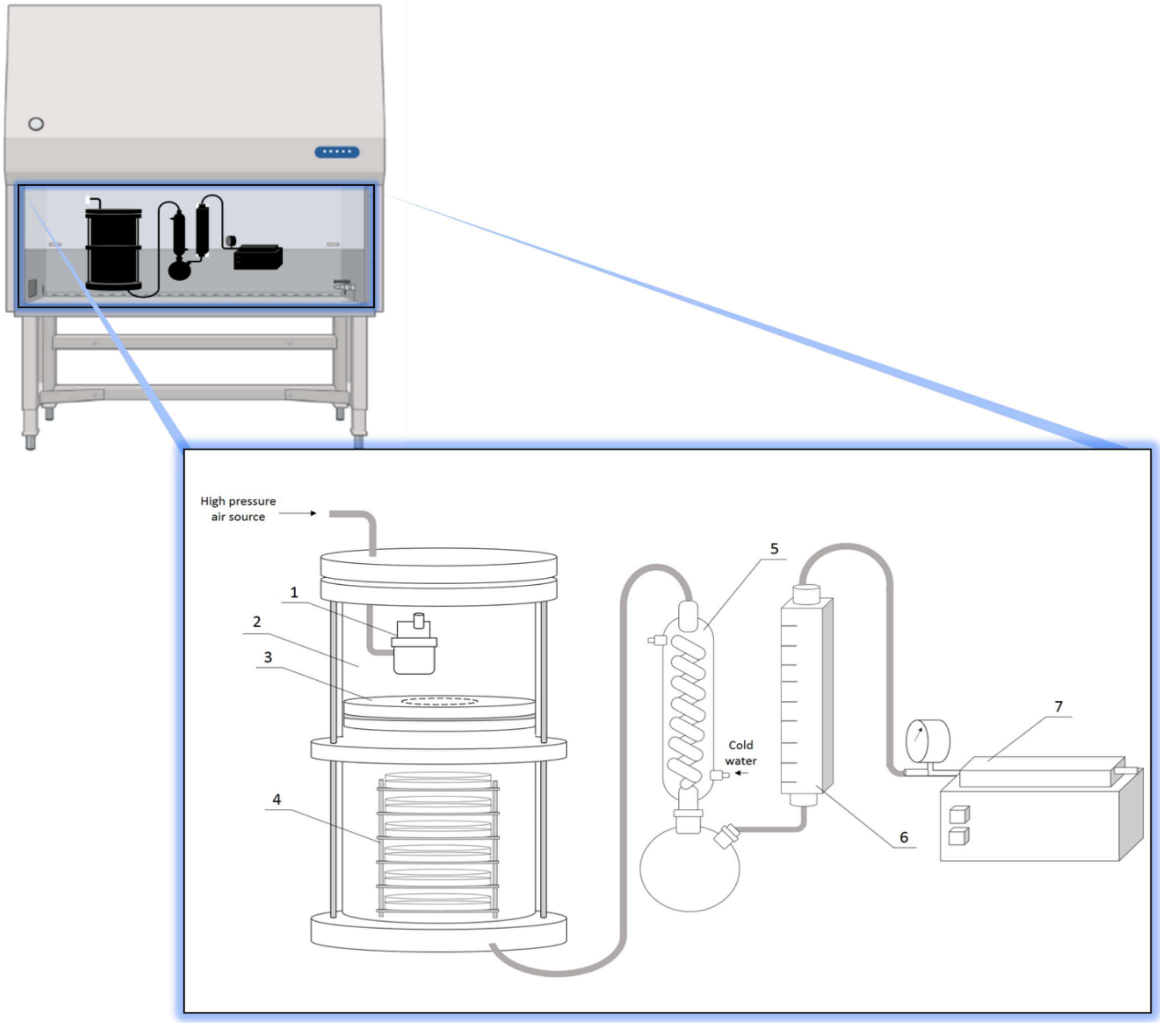


**Supplementary Figure 8**. Diagram of device used to measure the bacterial and viral filtration efficiency. 1 – nebulizer; 2 - aerosol chamber; 3 - insert with test filter; 4 - Petri dishes; 5 – condenser; 6 - calibrated flowmeter; 7 - vacuum pump.

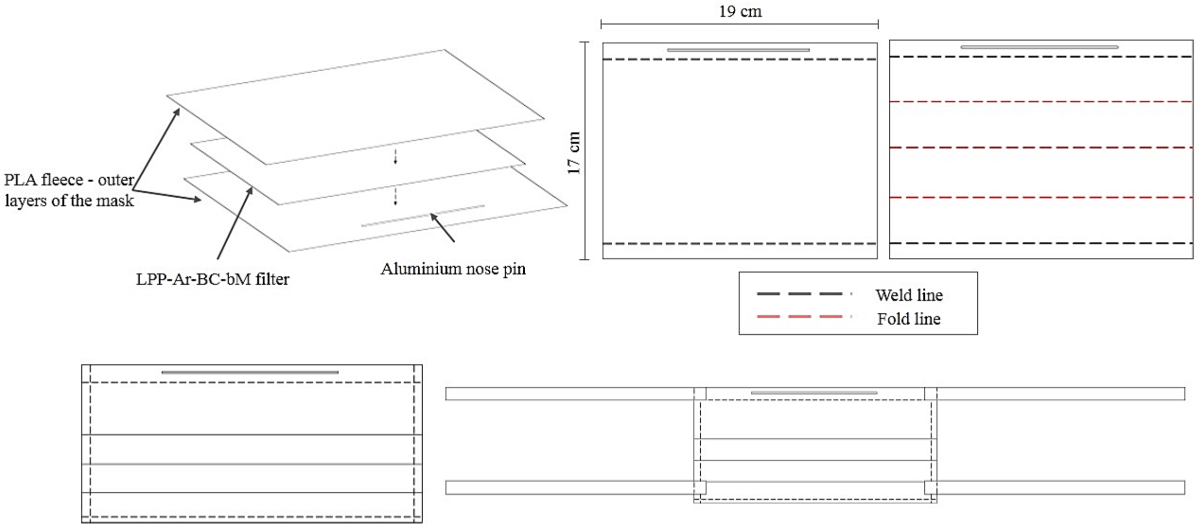


**Supplementary Figure 9.** Design of the NanoBioCell mask.

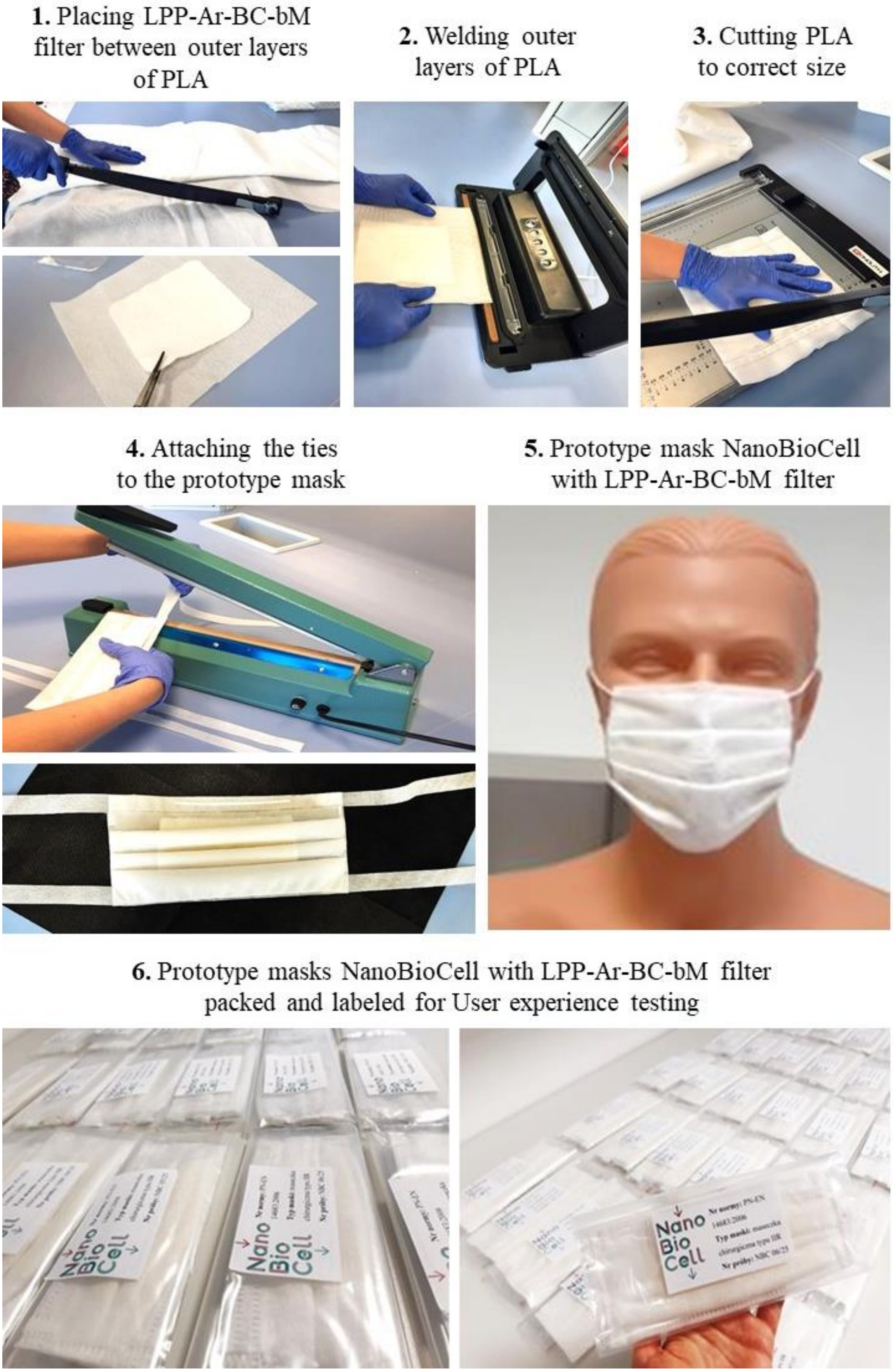


**Supplementary Figure 10.** Scheme of preparation of the NanoBioCell mask.
